# Combined blockade of VEGF, Angiopoietin-2, and PD1 reprograms glioblastoma endothelial cells into quasi-antigen-presenting cells

**DOI:** 10.1101/2022.09.03.506476

**Authors:** Zohreh Amoozgar, Jun Ren, Nancy Wang, Taylor P. Uccello, Patrik Andersson, Gino B. Ferraro, Shanmugarajan Krishnan, Pin-Ji Lei, Sonu Subudhi, Kosuke Kawaguchi, Rong En Tay, Igor L. Gomes-Santos, Peigen Huang, Hye-Jung Kim, Dai Fukumura, Rakesh K. Jain

## Abstract

Glioblastoma (GBM) remains a highly aggressive and uniformly fatal tumor, with a 5-year survival of <10%. Moreover, all randomized clinical trials with immune-checkpoint blockers have failed to date. Here we report that tumor endothelial cell (EC) dysfunction confers resistance to immunotherapy in preclinical GBM models. Anti-VEGF-therapy-induced vascular normalization is insufficient to fully restore the EC function and alleviate inflammatory edema induced by blocking programmed cell death protein 1 (PD1). Strikingly, concomitant blockade of angiopoietin 2 (Ang2), vascular endothelial growth factor (VEGF), and programmed cell death protein 1 (PD1) reprograms dysfunctional ECs to quasi-antigen presenting cells and upregulates receptors required for cytotoxic T lymphocyte entry into the tumor. Blocking VEGF, Ang2, and PD1 induces durable anti-tumor response while controlling edema. Upregulation of the transcription factor T-bet is necessary for generating resident memory T cells elicited by this combination therapy. Moreover, a human GBM organoid model formed from human cancer cells, ECs, CD14 and CD8 cells recapitulated these findings. Our study reveals the role of Ang2 in resistance to VEGF and/or PD1-blockade and provides a compelling rationale for clinical evaluation of this combination in GBM patients.

## Introduction

Glioblastoma (GBM) is a universally fatal brain tumor with a 5-year survival benefit of <10% from the current standard of care [surgical resection, radiation (RT), chemotherapy, anti-vascular endothelial growth factor (αVEGF) antiangiogenic therapy, and tumor treating fields] ^1^. Moreover, all randomized phase II and III trials using immune checkpoint blockers (ICBs) in GBM patients have failed to date ^1–3^. The limited efficacy of ICBs is partly due to the heterogeneity of transcriptional tumor states ^4^ and formidable barriers that the GBM tumor microenvironment (TME) creates to evade immune surveillance, block infiltration and function of antitumor immune cells and resist ICB therapies ^5–7^. GBM cancer cells are highly proliferative and invasive with a low neoantigen load and reduced MHC-I antigen presentation, allowing cancer cells to evade immune surveillance. Furthermore, GBM tumor vessels are morphologically and functionally abnormal, creating a hypoxic, acidic and edematous environment ^6,8–10^. The vascular abnormalities limit the delivery of drugs and infiltration of antitumor immune cells such as cytotoxic T lymphocytes (CTLs) into the tumor ^11,12^. In addition, the luminal surfaces of malformed vessels express low levels of endothelial cell adhesion molecules (ECAMs) on endothelial cells (ECs) ^6,8^ that are required for CTL homing to tumors, thereby limiting antitumor responses. ICB efficacy depends on sufficient CTL infiltration and activation ^6,8^. GBMs exclude CTLs from the TME and thus, are known to be immunologically “cold,” while pro-tumor immune cells including T regulatory cells (Tregs) and immunosuppressive macrophages (M2-like) infiltrate and expand in the GBM TME ^6,8,13–15^. Furthermore, the limited CTLs sequestered at the tumor margin are dysfunctional, exhausted, and express high levels of inhibitory receptors – including the programmed cell death receptor 1 (PD1) ^16–17^. Collectively, these inherent GBM features constitute a strongly immunosuppressive TME that mediates treatment resistance in GBM patients.

The ECs lining blood vessels form a barrier between circulating immune cells and parenchymal tissue. ECs in GBM are known to be dysfunctional ^6,8,10^. We and others have shown that normalizing vessels with antiangiogenic agents can improve multiple physiological features of the tumor, such as oxygenation, perfusion, and expression of ECAMs, e.g., ICAM1, VCAM1, and E-selectin ^2,6,8^, that facilitate adhesion and extravasation of leukocytes. Thus, vascular normalization contributes to significantly increased infiltration of endogenous CTLs ^18,19^ and adoptively transferred CAR-T cells^20^. However, CTL accumulation in the tumor bed is necessary but not sufficient for the therapeutic response, as seen in poor therapeutic outcomes in patients with inflamed tumors (e.g., colorectal, lung adenocarcinoma) ^21^. Sustained activation of CTLs by antigen-presenting cells is a key step for effective antitumor responses. Here, we propose that reprogramming the dysfunctional GBM ECs may help realize this goal by i) upregulating receptors on ECs that promote CTL trafficking, and ii) presenting tumor antigens to CTLs by ECs resulting in generation of an active tumor-immune niche in GBM.

Previous studies from our group and others have shown that antiangiogenic agents inhibiting VEGF signaling can transiently restore normal-like vascular function ^2,6^. However, tumor vessels become abnormal when the window of normalization closes via activation of Tie2 signaling in murine models of GBM as well as in GBM patients ^22,23^. Indeed, blocking both Angiopoietin2 (Ang2) and VEGF can extend the normalization window and increase survival of GBM-bearing mice ^24,25^. Provocatively, Ang2 levels also correlate with poor efficacy of αPD1 in melanoma patients ^26^. To this end, we used a bispecific antibody that blocks both Ang2 and VEGF (A2V) ^24^ with an αPD1 antibody in three preclinical models of GBM: GL261-MGH ^15^, a variant of GL261 ^24^, which is highly vascularized, but necrotic, invasive, and recapitulates resistance to αPD1 therapy seen in patients; CT2A for its relatively low neoantigen load and antigen presentation deficiency ^27–29^; and 005GSC that mimics stem-like features of GBM in patients ^14,15,30^. Here, we demonstrate that the concurrent blockade of VEGF, Ang2 and PD1 enables antigen presentation by ECs to re-activate anti-tumor lymphocytes (tumor specific CTLs as well as natural killer (NK) cells) and subsequent formation of memory T cells, resulting in increased anti-tumor immune response. Similarly, in *in vitro* human tumoroid model, A2V along with αPD1 treatment resulted in increased interferon gamma (IFNɣ) driven cancer cell control, dependent on MHC class I antigen presentation. Our study provides mechanistic insights into the resistance to anti-VEGF agents and immunotherapy in GBM, and supports clinical evaluation of concomitant blockade of VEGF, Ang2, and PD1.

## Results

### A2V overcomes resistance to αPD1 in murine GBM tumor models

To test if αVEGF therapy can overcome therapeutic resistance to αPD1 treatment, we treated tumor-bearing mice with B20, a monoclonal antibody against murine VEGF at the standard pre-clinical dose (6 doses of 5 mg/kg, bi-weekly, i.p.) alongside αPD1 (6 doses of 250 µg/dose, i.v. bi-weekly). We first tested our hypothesis using GL261-MGH model ^15^. Mice bearing size-matched GL261-MGH tumors (2 mm^3^) were treated with an isotype control antibody (6 doses of 5 mg/kg, bi-weekly, i.p.), B20 (5 mg/kg, twice a week), αPD1 (250 µg/dose), or B20+αPD1 (Fig. 1a). While αPD1 monotherapy did not enhance survival of mice, the combination of B20 with αPD1 modestly improved median overall survival from 20 days to 25 days, but without any durable response (Fig. 1b).

**Figure 1.**
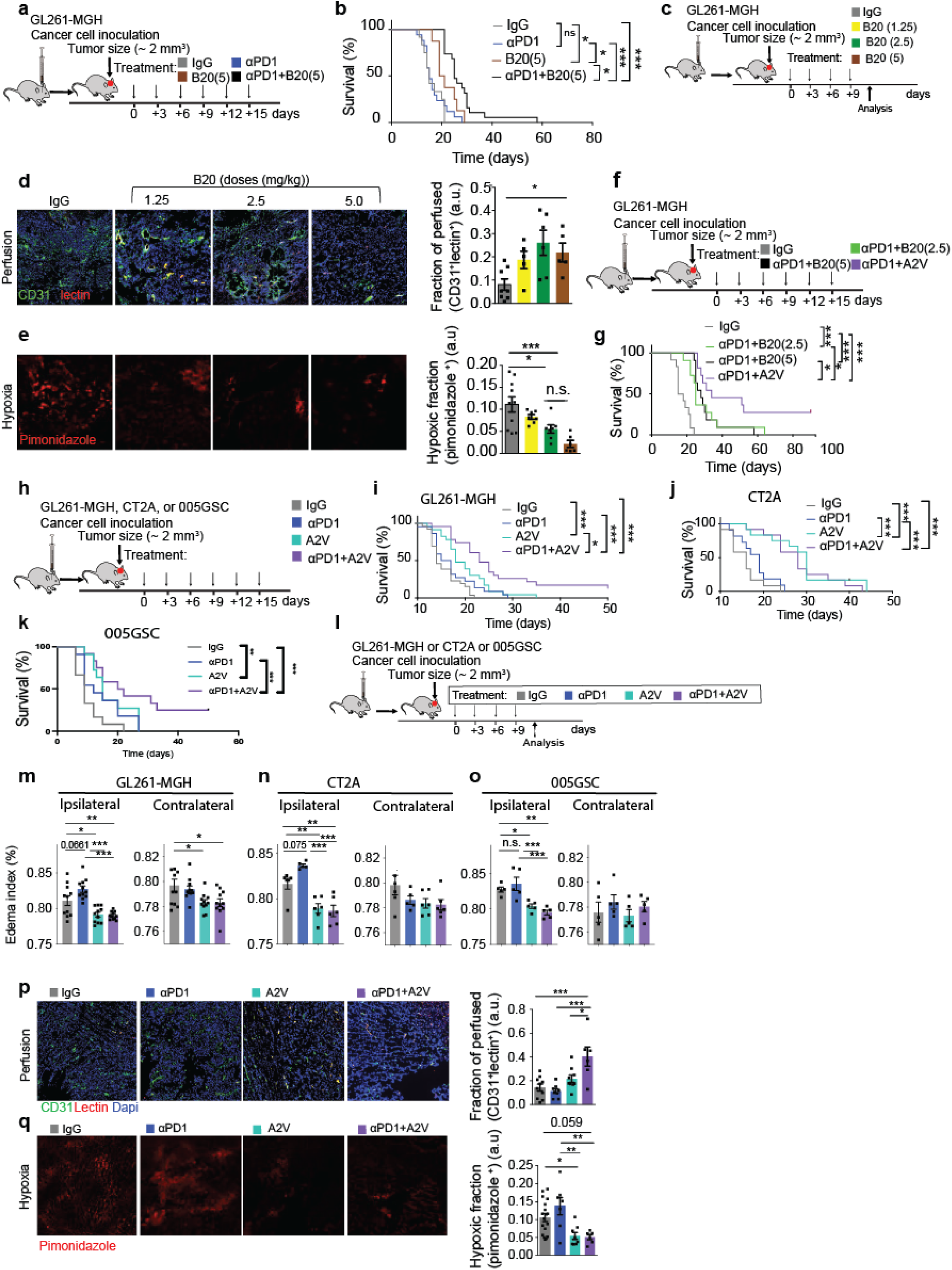
Resistance to αVEGF and αPD1 therapies is mitigated by simultaneously blocking Ang-2. **a)** Experimental design to evaluate the effect of IgG (isotype control), αPD1, αVEGF, and αPD1+αVEGF on the survival of GL261-MGH GBM-bearing mice (representative of two independent repeats of survival assessment). **b)** Kaplan-Meier survival curves. Median survival for GL261-MGH [IgG 250 μg/mice biweekly (n=12, 15 days), αPD1 250 μg/mice biweekly (n=17, 15 days), B20 5 mg/kg biweekly (n=8, 20 days), αPD1+B20 (n=19, 25 days) *p* <.0001]. **c)** Experimental design to evaluate the effect of dosing regimen of B20 on vascular normalization in mice. **d)** Measurement of perfusion in GL261-MGH tumors treated with B20. ***left***, Representative immunohistochemistry images of endothelial cells (CD31, green), perfused vessels (lectin, red) and nucleus, (DAPI, blue). ***right***, Quantification of perfusion treated with B20 1.25 mg/kg (n=5), 2.5 mg/kg (n=6), 5 mg/kg (n=5) or IgG (250 µg/mouse, n=8), *P*=0.0134, by ordinary one-way ANOVA test and corrected for multiple comparisons using the Tukey adjustment. For between group analysis post-Tukey, Statistical significance is shown as \**p* < 0.05, \*\**p* < 0.01, \*\*\**p* < 0.001, \*\*\*\**p* < 0.0001. **e)** Measurement of hypoxia in GL261-MGH tumors treated with B20 1.25 mg/kg (n=7), 2.5 mg/kg (n=7), 5 mg/kg (n=6) or IgG (250 µg/mouse, n=10). ***left***, Representative immunohistochemical images of pimonidazole staining (red, images demonstrate hypoxic area). ***right***, quantification of hypoxic area *P*=0.004 by ordinary one-way ANOVA test and were corrected for multiple comparisons using the Tukey adjustment. For between group analysis post-Tukey, Statistical significance is shown as \**p* < 0.05, \*\**p* < 0.01, \*\*\**p* < 0.001, \*\*\*\**p* < 0.0001. **f)** Experimental design to evaluate survival of GL261-MGH GBM-bearing mice after IgG, αPD1+B20 (2.5 mg/kg), αPD1+B20 (5 mg/kg), and αPD1+A2V treatments. **g)** Kaplan-Meier survival curves of the experiment in (**f**). Animal number and median survival: IgG (n=12, 16.5 days), αPD1+B20 (2.5) (n=11, 25 days), αPD1+B20 (5) (n=11, 28 days), αPD1+A2V (n=11, 34 days) *p* = <0.0001]. statistical significance is shown as \**p* < 0.05, \*\**p* < 0.01, \*\*\**p* < 0.001, \*\*\*\**p* < 0.0001. **h)** Schematic representation of experimental setup to evaluate survival of GL261-MGH, CT2A, 005GSC GBM-bearing mice after IgG, αPD1, A2V, and αPD1+A2V treatments. **i-k)** Kaplan-Meier survival curves. Median survival for GL261-MGH [IgG (n=26, 14 days), αPD1 (n=22, 16 days), A2V (n=23, 18days), αPD1+A2V (n=23, 23 days), *p* = <0.0001; and 3 out of 23 mice were tumor free]; CT2A [IgG (n=12, 16 days), αPD1(2.5) (n=11, 19 days), A2V (n=12, 30 days), αPD1+A2V (n=12, 28 days), *p* <0.0001]; 005GSC [IgG (n=12), 9 days), αPD1 (n=11, 12 days), A2V (n=11, 15 days), αPD1+A2V (n=12, 21 days), *p* = <0.0004; and 2 out of 12 mice were tumor free]; Statistical significance is shown as \**p* < 0.05, \*\**p* < 0.01, \*\*\**p* < 0.001, \*\*\*\**p* < 0.0001. **l)** Schematic of treatment-schedule/protocol used in **m-q**. **m)** Edema measurement in ipsilateral (side of the tumor, *p* < 0.0001) and contralateral brain hemispheres of GL261-MGH (N=11, *p*=0.0093). **n-o**) A2V and αPD1+A2V ipsilateral brain hemispheres in CT2A [IgG (n=6), αPD1 (n=5), A2V (n=6), αPD1+A2V (n=6), p < 0.0001]; and 005GSC (n=5, *p*=0.0002). Data presented are mean ± SEM. For between-group analysis post-Tukey, Statistical significance is shown as \**p* < 0.05, \*\**p* < 0.01, \*\*\**p* < 0.001, \*\*\*\**p* < 0.0001. **p)** Measurement of perfusion in GL261-MGH tumors treated as in **l**. Representative immunohistochemistry images of endothelial cells (green), perfusion (red) and nucleus (blue) (***left***) and quantification (***right***). Multiple comparisons were performed using the Tukey adjustment. For between-group analysis post-Tukey, [IgG (n=10), αPD1 (n=8), A2V (n=9), αPD1+A2V (n=6), *p*=0.0002] Statistical significance is shown as \**p* < 0.05, \*\**p* < 0.01, \*\*\**p* < 0.001, \*\*\*\**p* < 0.0001. **q)** Measurement of hypoxia in GL261-MGH tumors treated as in **l**. Representative immunohistochemical images of hypoxia staining (***left***) and quantification (***right***). Multiple comparisons were performed using the Tukey adjustment. [IgG (n=18), αPD1 (n=7), A2V (n=8), αPD1+A2V (n=6), *p*=0.0012] and statistical significance is shown as \**p* < 0.05, \*\**p* < 0.01, \*\*\**p* < 0.001, \*\*\*\**p* < 0.0001.

We have previously demonstrated in breast cancer ^19^ that lowering the dose of αVEGF extends the window of vascular normalization and consequently increases the delivery of antitumor immune cells and drugs to the tumors. Therefore, we sought to determine whether lower doses of B20 below the standard pre-clinical dose could normalize the vessels (Fig. 1c) and control GBM (Supplementary Fig. 1-2). Whereas B20 at 1.25 mg/kg was ineffective, 2.5 mg/kg increased vessel perfusion and reduced hypoxia (Fig. 1d-e) to a similar extent as the standard dose of 5 mg/kg. Similar to the standard dose (5 mg/kg), a lower dose of B20 (2.5 mg/kg) in combination with αPD1 was not sufficient to generate a durable response (Fig. 1f-g). These data indicate that in GL261-MGH, judicious dosing of B20 with αPD1 results in the similar therapeutic outcome as the standard dosing regimen.

We found that B20 at standard and reduced doses in combination with αPD1 extended the survival of GBM bearing mice only modestly. By contrast, αPD1 (250 µg/dose) + A2V (125 µg/dose) not only provided a superior survival benefit, but also completely eradicated tumors in 3 out of 23 mice (Fig.1g). We next investigated if other antiangiogenic blockers targeting pathways beyond VEGF, in combination with αPD1, could produce a similar antitumor response as αPD1+A2V. We selected regorafenib, a dual blockade of VEGFR2-Tie2 that showed promising results in a phase II GBM clinical trial but failed in randomized phase III trial ^31,32^. We observed that αPD1+A2V provided a superior survival benefit compared to regorafenib+αPD1 (Supplementary Fig. 3). Encouraged by our αPD1+A2V findings, we evaluated tumor growth and survival benefit following 6 doses of i) control: IgG2a (250 µg/dose) + IgG1 (MopC21, 125 µg/dose), ii) αPD1 (250 µg/dose), iii) A2V (125 µg/dose), and iv) αPD1+A2V (250 µg/dose and 125 µg/dose, respectively) bi-weekly (Fig. 1h) in all three GBM models. A2V improved survival compared to control and αPD1 in all three tumor models. CT2A was especially sensitive to A2V, as A2V alone doubled median survival of mice compared to control IgG. However, adding αPD1 to Α2V did not improve survival beyond A2V alone (Fig. 1j), presumably because of antigen presentation deficiency and mesenchymal differentiation in CT2A ^29^. By contrast, dual therapy with αPD1+A2V led to a significant increase in survival, including eradication of GL261-MGH and 005GSC in a fraction of mice (∼20%) (Fig. 1i, k).

Since we observed different degrees of therapeutic outcomes of αPD1+A2V in GL261-MGH, 005GSC and CT2A models and that the primary target of A2V is ECs ^24,25^, we next investigated the extent of vascular normalization under each therapy. Edema is a major adverse consequence of abnormal vessels in GBM and can be controlled by vascular normalization ^23,33,34^. However, αPD1 can exacerbate edema in GBM in mice as well as patients ^13,35^. Therefore, we evaluated edema in GL261-MGH, CT2A, and 005GSC tumor models following control (IgG), monotherapies (αPD1 or A2V), or dual therapy (αPD1+A2V) (Fig. 1l-o). Both A2V and αPD1+A2V therapies reduced edema in the ipsilateral and contralateral brain hemispheres in mice bearing GL261-MGH tumors (Fig. 1m). However, the extent of edema control in CT2A and 005GSC was limited to the ipsilateral brain hemisphere (Fig. 1n-o). Given that GL261-MGH is refractory to αPD1, similar to GBM in patients, we selected this model for further analysis using functional, genetic and proteomic approaches. As anticipated, analysis of vascular function showed increased vessel perfusion and reduced hypoxia in GL261-MGH tumors treated with αPD1+A2V compared to the other three regimens (Fig. 1p-q).

### αPD1+A2V converts dysfunctional GBM ECs to quasi-professional antigen-presenting cells

To validate our findings in the context of human cells, we exposed human ECs and primary monocytes (CD14+) to human tumor conditioned media (U87 cell line) and observed that VEGFR2 expression was reduced on both cell populations following αPD1+A2V therapy (Fig. 2a, b). Additionally, combination therapy resulted in a decrease in Tie2 expression in ECs (Fig. 2c). Transcriptomic analysis of ECs isolated from murine GBM showed that genes expressed in ECs after αPD1+A2V therapy cluster separately from those expressed with monotherapies (Fig. 2d-e). The primary determinants of PC1 are vascular-specific genes that track the successful morphological and functional transition of tumor vessels toward a normalized state (Supplementary Fig. 4a-c). Together these data reveal a potential mechanism by which abnormal angiogenesis is suppressed following αPD1+A2V treatment.

**Figure 2.**
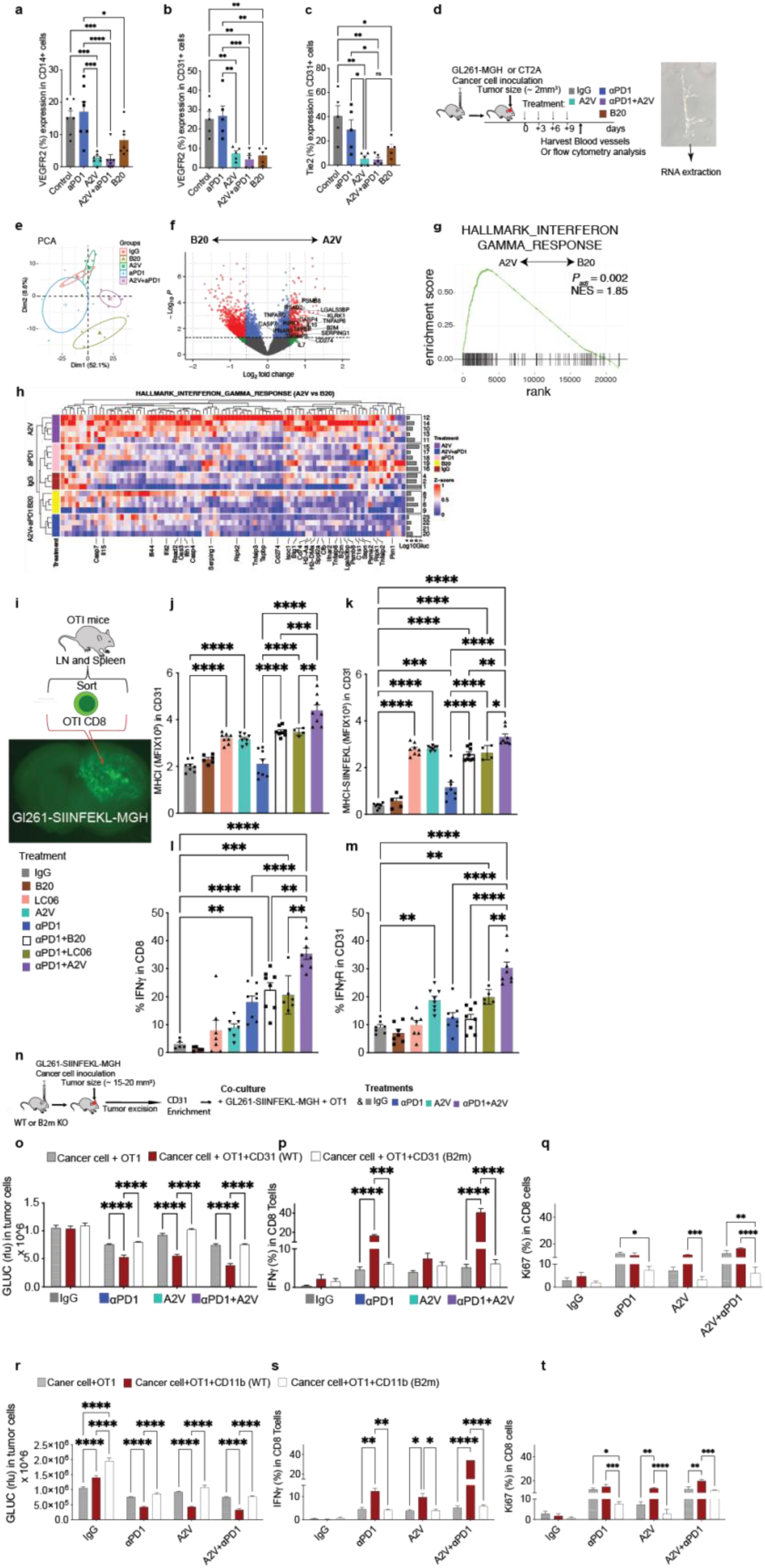
αPD1+A2V transforms the phenotype and the function of endothelial cells. **a-b**) VEGFR2 expression on (**a**) human CD14+ monocytes and (**b**) human CD31+ ECs exposed to tumor conditioned media and treated with Control, αPD1, A2V, αPD1+A2V, or B20. **c)** Tie2 expression on human CD31+ ECs exposed to tumor conditioned media and treated with Control, αPD1, A2V, αPD1+A2V, or B20. **d)** Schematic representation of treatment schedule/protocol, harvesting tumor blood vessels, and analysis of endothelial cells (GL261-MGH RNA sequencing and GL261-MGH and CT2A flow cytometry). **e)** Principal components analysis showing principal component 1 (PC1) and PC2 for RNA-seq data from tumor vessels (GL261-MGH) treated with IgG (n=3, orange), αPD1 (n=5, cyan), B20 (n=4, gold), A2V (n=4, green), αPD1+A2V (n=4, purple). The ellipses around each group represents 95% confidence level. **f)** Volcano plot showing differentially expressed genes (DEGs) between B20 and A2V. **g**). Gene set enrichment analysis (GSEA) showing enrichment of Interferon pathways in A2V treatment versus B20. **h)** Heatmap showing Hallmark Interferon Gamma Pathway related genes that contributed to enrichment of the gene set in A2V in comparison to B20. **i)** Schematic showing preparation of slice culture from GL261-SIINFEKL-MGH cells, which preserves the TME, including their structural and cellular elements (e.g., myeloid cells). **j)** MHCI [Mean Fluorescent Intensity (MFI)] **k)** and MHCI (SIINFEKL) (MFI) expression on ECs. **l)** IFNγ production by CD8 T cells **m)** and IFNγR expression on ECs after therapy as indicated in schematic (**i**). Data presented are mean ± SEM. Multiple comparisons using the Tukey adjustment. For between-group analysis post-Tukey, Statistical significance is shown as \**p* < 0.05, \*\**p* < 0.01, \*\*\**p* < 0.001, \*\*\*\**p* < 0.0001. **n)** Schematic of *in vitro* co-culture assay. GL261-SIINFEKL-MGH cells are orthotopically implanted in mice. Tumors are excised when they reach a size of 15-20 mm^3^. To enrich endothelial cells, whole blood vessels are harvested using ficoll gradient separation and then dissociated into single cells using 5 min accutase ® exposure at room temperature. WT CD8 T cells and OTI cells were enriched from the spleen using Stem Cell Technologies negative selection kit to a purity of 94-98%. CD11b cells were isolated from the bone marrow. Cells were co-cultured and treated as described in this panel. **o, r**) Production of GLUC by cancer cells as a measure of cancer cell viability after therapy, (**p, s**) IFNγ production by OTI cells in the co-culture of cancer cells (GL261-SIINFEKL-MGH) with ECs (CD31), harvested from WT versus *B2m*-KO mice. (**q, t**) Ki67 expression by OT-1 cells in the co-culture. Data presented are mean ± SEM. Multiple comparisons using the Tukey adjustment. For between group analysis post-Tukey, stars were assigned as \**p* < 0.05, \*\**p* < 0.01, \*\*\**p* < 0.001.

Although previous studies showed that targeting Ang2 and VEGF can enhance vascular normalization to a similar extent in melanoma ^26^, our study revealed that αPD1+A2V increased survival more than αPD1+B20 in GBM (Fig. 1g). To delve deeper into the underlying mechanisms involved in this differential response, we compared the differences in MSigDB hallmark gene sets in tumor-associated ECs isolated from mice treated with A2V versus B20. Interestingly, we found that the expression of genes involved in antigen processing and presentation function of APCs was significantly higher in A2V treated ECs in comparison to B20 treatment ones (Fig. 2f). These differentially expressed genes include: i) *TAPBP* and *B2m* that are critical for the peptide transport and MHC assembly ^36–38^, ii) *PSMB8*, a subunit of the (immuno) proteasome which degrades proteins into the peptides presented on MHC I molecules ^39^, iii) *RSAD2* known to be critical for DC maturation ^40^ and iv) *CD274* encoding PDL1 that is known to be upregulated secondary to IFNγ stimulation ^41^. Additionally, we found that the response to IFNɣ signaling was higher in A2V treated ECs compared to B20 treatment (Fig. 2g, h). These data collectively suggest that A2V not only contributes to the normalization of vasculature supporting immune cell infiltration into the tumor but may also promote T cell function potentially through enhanced presentation of tumor associated antigens (TAAs) and priming towards an IFNɣ primed subset of T cells by ECs. The upregulation of PDL1 on ECs after A2V treatment, on the other hand, suggests that adding αPD1 may boost the efficacy of A2V treatment by enhancing CD8 T cell function.

To delve more deeply, we ran CIBERSORT using the brain-specific gene signatures, and identified that endothelial-like and, fibroblast-like cells were enriched in our datasets, suggesting that the majority of the genes in our isolated vessel RNA-Seq data were derived from brain vasculatures. Of note, the macrophage-like features are also enriched in our samples (Supplementary Fig. 4d-g). However, we did not observe a significant increase in the macrophage population in the αPD1+A2V group in comparison to other samples (Supplementary Fig. 4d-g). This indicates that the IFN signature that we observed upon GSEA analysis was not driven by higher macrophage-like features, suggesting that this change in gene expression is treatment-driven. Finally, consistent with reduced tumor hypoxia following vascular normalization, GSEA revealed significant downregulation of the hypoxia-related pathways in αPD1+A2V treated tumors compared to IgG controls (Supplementary Fig. 5a). To facilitate comparison of IFNγ pathway activation between A2V and B20 treatment, we have included plots of DESeq2 size-factor normalized counts for key genes in the Supplementary Materials (Supplementary Fig. 5b). Genes including *Cd274, B2m, Stat1, and Irf1* show normalized count values ranging from several hundred to over 5,000 counts across A2V replicates, well above the technical background threshold, supporting the biological validity of the endothelial-immune crosstalk.

Based on these data suggesting increased expression of genes for antigen presentation machinery by ECs after A2V treatment, we posited that the reprogrammed GBM ECs may have gained an ability to present antigens to CD8 T cells. To test this hypothesis, we developed an antigen-specific model by integrating the SIINFEKL part of ovalbumin (OVA) into the GL261-MGH genome to establish a GL261-MGH-OVA model. OVA peptide SIINFEKL was efficiently presented in the context of H-2K^b^ in this model (Supplementary Fig. 6). Moreover, the immunogenicity of the GL261-MGH was not altered by SIINFEKL expression, as GL261-SIINFEKL-MGH and GL261-MGH showed similar tumor growth in wildtype (WT) C57BL6 mice (Supplementary Fig. 7).

To test the ability of ECs to present tumor-associated antigens (TAA) to CD8 T cells, we adopted an organotypic co-culture model that remains viable for more than two weeks ^42,43^. Similar to *in vivo* studies, we implanted the brain slice cultures with GL261-SIINFEKL-MGH cells (Fig. 2i), randomized the cultures based on tumor size and then treated with 1) control: IgG2a (10 μg/mL) +MOPC21 (10 μg/mL), 2) αPD1 (10 μg/mL), 3) B20 (10 μg/mL), 4) LCO6 (anti-Ang2 antibody) (10 μg/mL), 5) A2V (10 μg/mL), 6) αPD1+B20 (10 μg/mL+10 μg/mL), 7) αPD1+LCO6 (10 μg/mL+10 μg/mL) and 8) αPD1+A2V (10 μg/mL +10 μg/mL) in the presence of OTI CD8 T cells (Fig. 2i). We measured the expression of MHC class I on ECs, the proliferation of CD8 T cells, and cancer cell killing ^44^. These analyses were designed to capture key events in the cycle of antigen presentation, including tumor-associated OVA peptide loading onto MHC class I molecules and transport to the surface, TCR recognition of MHC class I/peptide and activation and differentiation of CD8 T cells into effector cytotoxic T lymphocytes (CTL), and CTL-mediated killing of target cells that express cognate MHC class I peptide complexes.

αPD1+A2V therapy induced the highest expression of MHC class I on ECs among these treatment conditions (Fig. 2j-k), which correlated with the highest IFNγ response by OT-I CD8 T cells (Fig. 2l). Responsiveness to IFNγ by stromal cells has been shown to be insufficient for IFNγ-induced tumor regression, whereas responsiveness of ECs to IFNγ is both necessary and sufficient ^45^. Moreover, preserved IFNγ receptor (IFNγR) signaling is required for CTL killing of cancer cells in GBM and other solid tumors ^46^. Analysis of IFNγR expression by ECs under each therapeutic setting revealed that αPD1+A2V therapy induced significantly higher expression of IFNγR on ECs compared to other treatment conditions (Fig. 2m). These data indicate that increased antigen presentation by ECs resulting from therapy-mediated normalization of ECs may contribute to the tumor regression and enhanced survival outcome because of enhanced CD8 T cell activity (Fig. 1i-k).

After taking a more stringent measure by including two arms (with/without αPD1) of bispecific A2V or [LCO6 (αAng2) or B20 (αVEGF)] alone, we observed that unlike B20 which had limited effect on the induction of MHC class I, LC06 significantly contributed to MHC class I upregulation in ECs (Fig. 2j). Overall, increased IFNγ expression was directly associated with αPD1 treatment as demonstrated by the significant difference between αPD1 vs control or A2V treatment groups (Fig. 2l).

To evaluate the relative contribution of ECs versus conventional antigen-presenting cells to the presentation of antigens to T cells, we developed multiple co-culture model systems composed of OT-1+ CD8 T cells with either ECs or CD11b+ cells. We sorted ECs and CD11b+ cells from GL261-SIINFEKL-MGH tumors in WT or *B2m* KO mice (Fig. 2n, Supplementary Fig. 8a). We treated cell co-cultures comprising GL261-SIINFEKL-MGH cells, OT-1 cells, and ECs or CD11b+ cells with 1) control: IgG2a + IgG1 (10 μg/mL+10 μg/mL), 2) αPD1 (10 μg/mL), 3) A2V (10 μg/mL), and 4) αPD1+A2V (10 μg/mL+10 μg/mL). These treatments did not alter cancer cell viability unless OT-I cells were present. Addition of WT ECs to the culture significantly reduced cancer cell viability after αPD1, A2V or αPD1+A2V treatment but this effect was lost when ECs were unable to present antigens to CD8 cells (*B2m*^-/-^ ECs) (Fig. 2o). Analysis of ECs from these culture systems showed that expression of MHC class I and MHC class I-SIINFEKL was at the highest level after αPD1+A2V therapy (Supplementary Fig. 8b-c). OT-I cells alone induce MHC class I expression on ECs, and this effect is doubled when both OT-I and CD11b⁺ cells are co-cultured with ECs (Supplementary Fig. 8b-c). Increased activation and proliferation of CD8 T cells was directly linked to the recognition of MHC class I on ECs or CD11b+ cells, since CD8 T cells co-cultured with ECs and CD11b+ cells from *Β2m* KO mice produced lower levels of IFNγ (Fig. 2p) and reduced proliferation (Ki67+) (Fig. 2q, Supplementary Fig. 8d-i). Interestingly, unlike ECs that did not actively induce tumor growth, inclusion of CD11b+ cells boosted tumor growth, particularly if they were unable to present antigen (∼1.7 fold). A2V therapy significantly reduced tumor control in the CD11b+/OT-1 cocultures (Fig. 2r) and elevated IFNɣ levels (Fig. 2s) but had marginal effect on proliferation compared to other treatments (Fig. 2t).

Moreover, we tested the response of OT-1 CD8 T cells in slice culture, where we generated brain slices from tumors grown in *B2m* KO mice or WT mice to test the interaction of OT-1 CD8 T cells either directly or indirectly using trans well culture (Supplementary Fig. 9). We cultured tumor-bearing slices in the top chamber and randomized based on the Gluc values measured in the media, after one or two days. To measure the impact of antigen presentation on T cell killing, we either added OT-1 cells on top of the slice culture in the top chamber (direct) or to the bottom chamber (indirect). We observed a strong tumor control when slice cultures were treated with αPD1+A2V and specifically when OT-1 cells were in direct contact with the tumors, suggesting the importance of antigen presentation and direct interaction with T cells (Supplementary Fig. 9a). Our data showed that the ability of T cells to kill cancer cells was significantly reduced in *B2m* KO mouse-derived slices, with impaired antigen presentation (Supplementary Fig. 9b). Additionally, the frequency, phenotype, and function of OT-1 cells that were directly exposed to cancer cells differed significantly from OT-1 cells that were added to the lower chamber (Supplementary Fig. 9c, d). OT-1 cells treated with αPD1+A2V and in direct contact with tumor slices were higher in frequency and anti-tumor phenotype marked by higher expression of CD44 and production of IFNɣ (Supplementary Fig. 9e-a10). The same phenotype was not observed in *B2m* KO slice culture, and specifically, OT-1 cells were lower in frequency regardless of direct or indirect contact with the slice cultures.

A2V is a cross-reactive antibody that recognizes and binds to both murine and human antigens. To evaluate if the antigen presentation mechanism observed in murine TME is conserved in human GBM, we developed and characterized four distinct human GBM tumoroids (Fig. 3a). These four models contained human GBM lines (U87, U251, U373, and SNB19) co-cultured with HUVECs, and CD14 and CD8 T cells isolated from patient PBMCs. GBM tumoroids were treated with 1) control: IgG2a (10 μg/mL), 2) αAng2 (10 μg/mL), 3) αVEGF (10 μg/mL), 4) αPD1 (10 μg/mL), 5) A2V (10 μg/mL), 6) αPD1+αAng2 (10 μg/mL+10 μg/mL), 7) αPD1+αVEGF (10 μg/mL+10 μg/mL), and 8) αPD1+A2V (10 μg/mL+10 μg/mL). In all four tumoroid models, dual combination of αPD1+A2V had the highest tumor control (Fig. 3b-e). Furthermore, combination treatment led to an increased inflammatory cytokine response (Fig. 3f, g) dependent on conventional MHC class I as the response was lost when *B2m* was depleted from the cancer cells (Fig. 3h, i).

**Figure 3.**
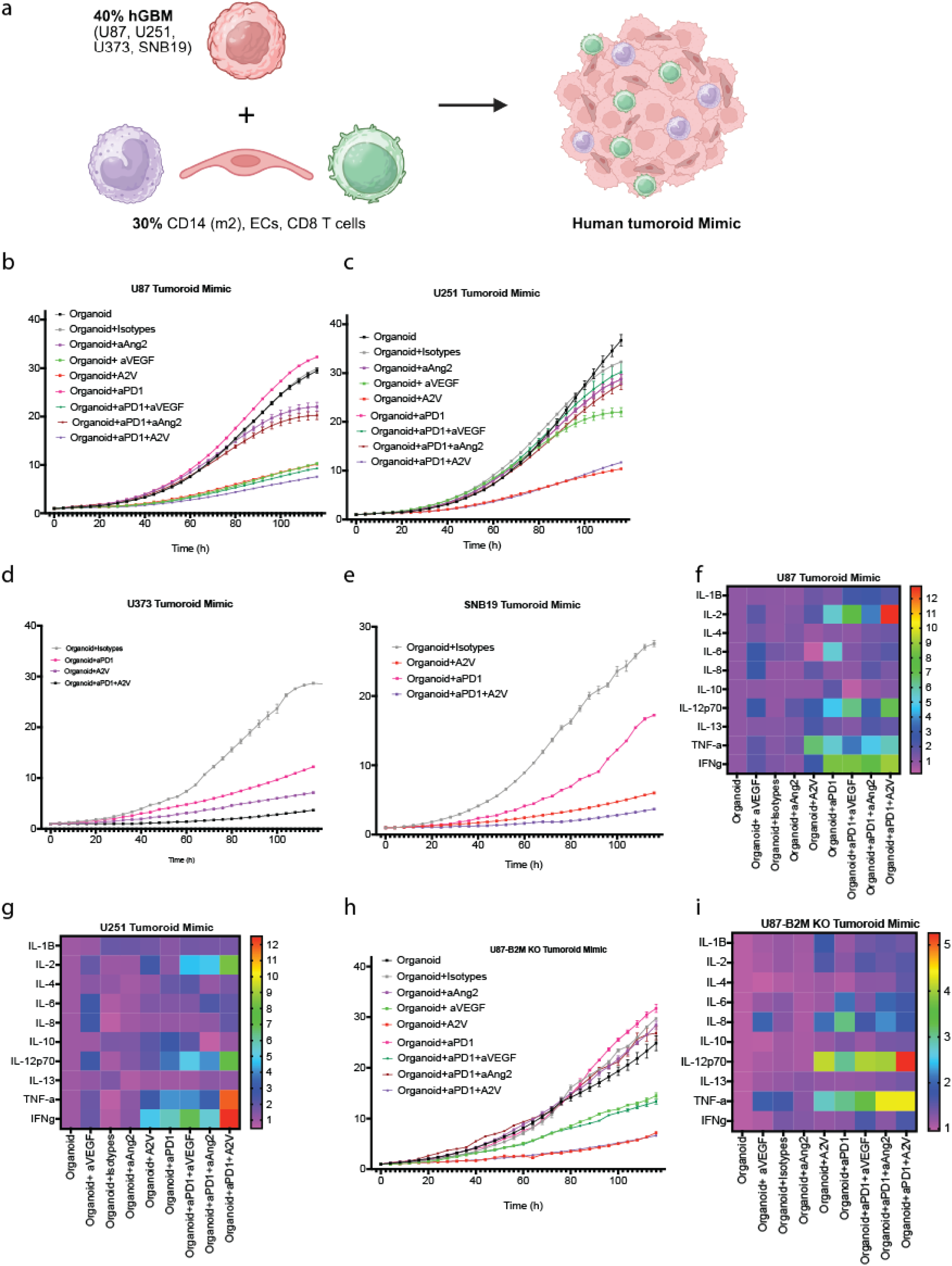
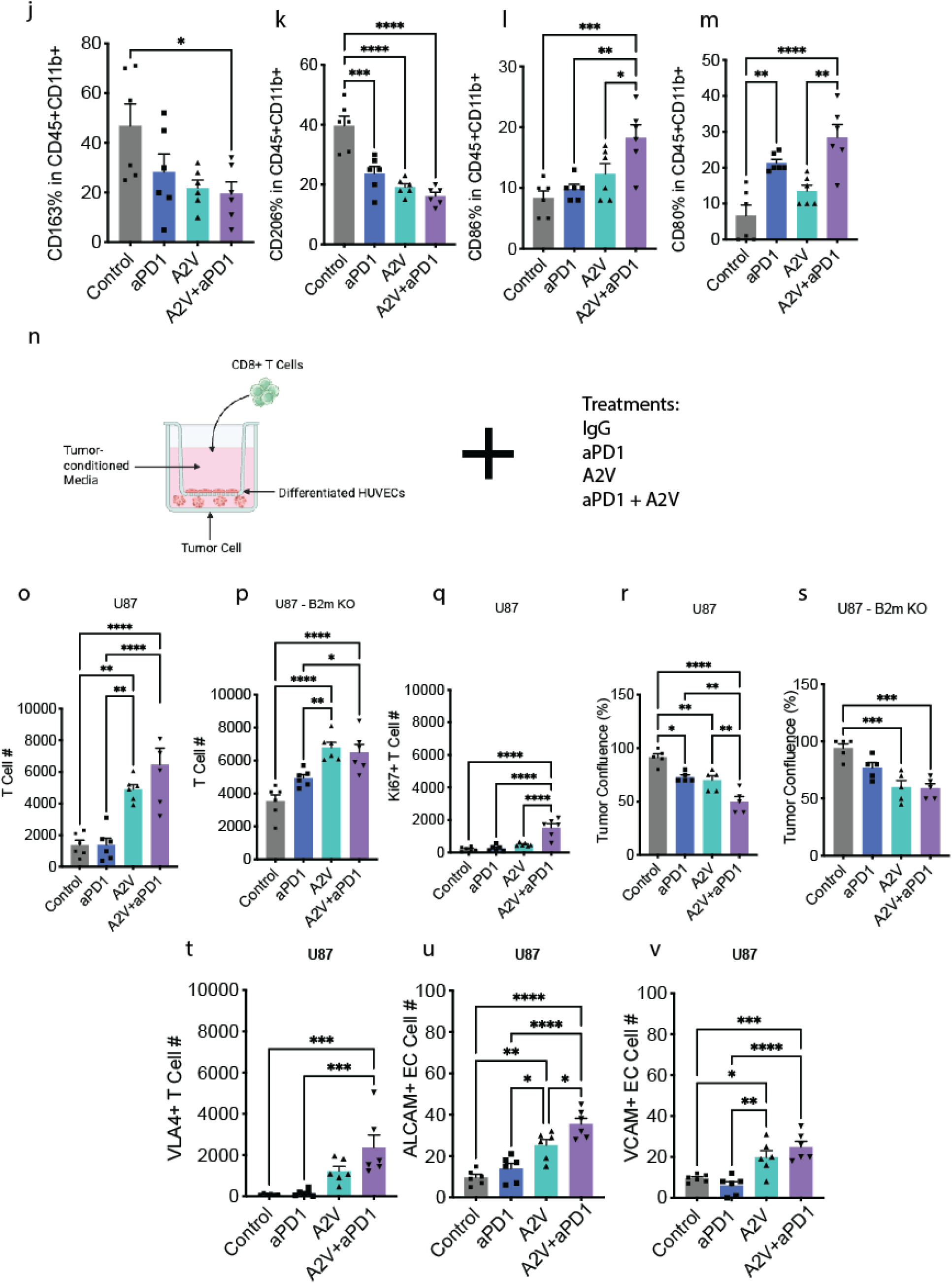
αPD1+A2V decreases cell viability in human GBM tumoroids and treatment stimulates IFNɣ in an MHC class I dependent manner. **a)** Schematic of tumoroid model. Human GBM cell lines (U87, U87-*B2M* KO, U251, U373, and SNB19) were cultured with differentiated human CD14 cells (M2- like), human ECs, and human CD8 T cells. When tumoroids formed, they were treated with IgG, αAng2, αVEGF, A2V, αPD1, αPD1+αVEGF, αPD1+αAng2, or αPD1+A2V. **b-e, h)** Cancer cell viability was assessed by Incucyte. **f, g, i)** Cytokine array for treated human tumoroids. **j-m)** Myeloid cell phenotyping by FACs. **n) Schematic of the** T cell extravasation set-up. Human GBM cell line (U87) or genetic variant with *B2M* KO (U87-*B2M* KO) were cultured with tumor conditioned media and HUVEC, treated with IgG, αPD1, A2V or αPD1+A2V, and cultured with CD8 T cells. **o-p)** T cell number in the bottom chamber was calculated. **q**) T cell Ki67 staining assessed by flow cytometry. **r, s**). Cancer cell confluency was measured by flow cytometry. **t-v**) VLA4+ T cells, ALCAM+ ECs, and VCAM+ ECs assessed by flow cytometry. n = 6

We then phenotyped the CD14 myeloid cells from the tumoroids following the treatment and found that αPD1+A2V led to a reduction in M2-like (CD136+, CD206+) myeloid cells and an increase in M1-like cells (CD86+, CD80+) (Fig. 3j-m).

The presence of ECAMs on ECs is an indicator of normalized vessels and facilitates adhesion and extravasation of leukocytes into tumors. We tested a series of ECAMs: peripheral lymph node addressin (PNAd) (Supplementary Fig. 10a-d), Intercellular Adhesion Molecule 1 (ICAM1), Vascular Cell Adhesion Molecule 1 (VCAM1), E-selectin and Activated Leukocyte Cell Adhesion Molecule (ALCAM)] that are known to contribute to CD8 T cell trafficking (Supplementary Fig. 10e-f). Among these adhesion molecules, ALCAM was highly expressed on ECs after αPD1+A2V therapy. In both GL261-MGH and CT2A models, ALCAM was significantly increased in A2V and αPD1+A2V groups (Supplementary Fig. 10g). However, there was no difference in ALCAM expression between A2V versus αPD1+A2V therapy in CT2A. Since ALCAM mediates leukocyte trafficking across the blood tumor barrier (BTB), this lack of ALCAM upregulation in CT2A may have contributed to the lack of therapeutic response in this model in contrast to GL261-MGH (Fig. 1i, j). Significant increase of *ALCAM* and *VCAM1* expression in ECs was also confirmed at mRNA level after αPD1+A2V therapy (Supplementary Fig. 10h-i).

We additionally tested the contribution of T cell trafficking using our human GBM cells in a dual-chamber experiment using differentiated HUVECs, tumor conditioned media with tumor antigens, human GBM cancer cells, and CD8 T cells (Fig. 3n). We first layered HUVECs on top of a 24-well chamber with cancer cells on the bottom and tumor conditioned media and antigen in between. We then added isolated CD8 T cells to the top chamber, treated co-cultures with 1) control: IgG2a (10 μg/mL), 2) αPD1 (10 μg/mL), 3) A2V (10 μg/mL), 4) αPD1+A2V (10 μg/mL+10 μg/mL) and measured T cell infiltration through the HUVECs and into the tumor layer (Fig. 3o, p). We found that while A2V as a monotherapy had an effect on T cell migration into the WT cancer cell layer, the combination of αPD1+A2V had the greatest effect. Similarly, we saw an increase in T cell migration into the *B2m* KO cancer cell layer with A2V monotherapy, indicating the HUVECs may serve as a source of antigen presentation to recruit CD8 T cells. However, in the *B2m* KO tumor co-culture, combined αPD1+A2V therapy was no better than A2V alone. Proliferative Ki67+ CD8 T cells were significantly increased in the αPD1+A2V treated group (Fig. 3q). Furthermore, αPD1+A2V therapy had the greatest impact in reducing cancer cell numbers and cancer cell confluency was not altered by the combination of αPD1+A2V in the *B2m* KO cancer cells, thus indicating the absence of MHC class I mediated antigen presentation on cancer cells, differentiated HUVECs provide an additional source of antigen presentation (Fig. 3r, s). In these co-cultures, we examined T and EC adhesion molecules’ expression and found that αPD1+A2V treatment resulted in increased VLA4 expression on T cells (Fig. 3t), as well as elevated ALCAM (Fig. 3u) and VCAM (Fig. 3v) on ECs.

### αPD1+A2V converts cold CTL-excluded GBM into inflamed tumors

We next analyzed whether αPD1+A2V therapy could improve lymphocyte trafficking into GBM as a result of their normalized vasculature (Fig. 1i-q) and transformed ECs (Fig. 2). We collected tumors from GL261-MGH and CT2A bearing mice after treatment with 4 i.v. doses of 1) control: IgG2a (250 µg/dose) + MOPC-21 (125 µg/dose), 2) αPD1 (250 µg/dose), 3) A2V (125 µg/dose), and 4) αPD1+A2V (250 µg/dose + 125 µg/dose) bi-weekly (Fig. 4a). GL261-MGH tumors were CTL-excluded, while CT2A tumors had CTL infiltration (Fig. 4b-c). Although treatment with αPD1 did not result in T cell re-localization (Fig. 4b), A2V and αPD1+A2V re-localized CTLs into the core of tumors in GL261-MGH tumor (Fig. 4b). In addition, total CTL numbers and the functional CTLs (*i.e.,* producing IFNγ and TNFα) were significantly increased after αPD1+A2V therapy compared with control or αPD1 monotherapy in GL261-MGH tumors (Fig. 4d-e). In CT2A tumors, A2V treatment increased functional CTLs compared to αPD1, without altering the overall number of CTLs (Fig. 4c, d). Moreover, αPD1+A2V therapy shifted the phenotype of CD8 T cells from exhausted to effector (Fig. 4e). Given that αPD1+A2V did not confer a survival advantage beyond A2V alone in CT2A tumors (Fig. 1j), we assessed whether differences in CTL localization and functional state distinguish these two GBM models. Consistent with granzyme B (GzmB) as a key effector of CD8 T cell cytotoxicity, CTLs infiltrating GL261-MGH tumors following αPD1+A2V exhibited significantly lower GzmB expression compared to CT2A, where CTLs displayed high basal levels of GzmB.

**Figure 4.**
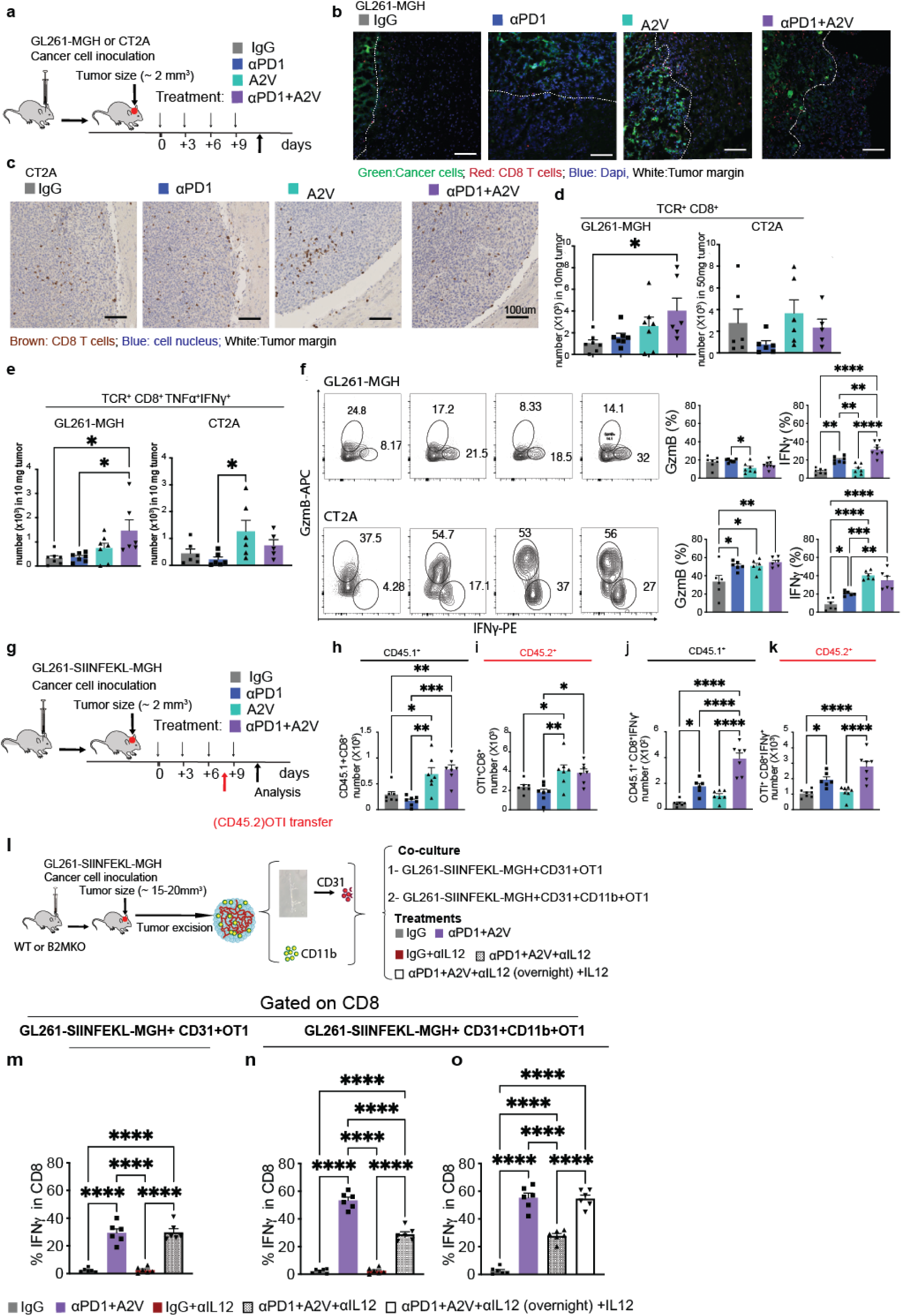
αPD1+A2V increases activated CTLs within GBMs. **a)** GL261-MGH and CT2A tumors were treated as in the schematic. **b-c)** Location of CTLs inside the GBM GL261-MGH [IgG (n=7), αPD1 (n=7), A2V (n=7), αPD1+A2V (n=6) *p* = <0.0410] (**b**); and CT2A [IgG (n=6), αPD1(n=6), A2V (n=6), αPD1+A2V (n=5), *p* = 0.0578] (**c**). **d-e**) Number of effector CTLs (**d**) which express cytokines (IFNγ^+^TNFα^+^) **(e**) inside the GBM GL261-MGH [IgG (n=7), αPD1 (n=7), A2V (n=7), αPD1+A2V (n=6), *p* = <0.0095]; and CT2A [IgG (n=6), αPD1 (n=6), A2V (n=6), αPD1+A2V (n=5), *p* = <0.074]. **f)** Inflammatory cytokines in CD8 T cells in GL261-MGH versus CT2A. ***Left***, FACS plots showing GzmB versus IFNγ production in CD8 T cells within GL261-MGH (**top**) compared with CT2A (**bottom**). ***Right***, Quantification of the production of GzmB versus IFNγ in CD8 CTLs in GL261-MGH (**top**) versus CT2A (**bottom**). **g)** Schematic of experimental design. GL261-SIINFEKL-MGH cells were orthotopically implanted in CD45.1 mice and, after tumors reached ∼2 mm^3^, mice were randomized into 4 treatment groups. After the third treatment, OTI cells were harvested and transferred into CD45.1 hosts. Mice received the fourth treatment and were then sacrificed, and tumors were analyzed for the presence of endogenous CD8 T cells and transferred antigen-specific CTLs [Vascular normalization enabled the trafficking of endogenous CTLs (CD45.1)]: (**h**) a combination of sequestered and systemic CTLs and (**i**) exogenous antigen-specific OTI (CD45.2). In both cases (**j-k**), αPD1 therapy increased the number of IFNγ producing CTLs. n=5-6. \**P*<0.05, \*\**P*<0.01, \*\*\**P*<0.001, \*\*\*\**P*<0.0001. **l**) Schematic representation of *in vitro* co-culture assay. GL261-SIINFEKL-MGH cells were orthotopically implanted in mouse brain. Tumors were excised and processed as in Supplementary Figure 10 and co-cultures were treated as shown in the supplementary figure 11a. **m-o**) Frequency of IFNγ+ CD8 T cell in (**m**) co-culture of cancer cells (GL261-SIINFEKL-MGH) with ECs (CD31) and ECs and antigen specific CD8 T cells (OTI) and (**n-o**) co-culture of cancer cells (GL261-SIINFEKL-MGH) with ECs (CD31), classical antigen presenting cells (CD11b), and antigen specific CD8 T cells (OTI).)

Since environmental cues orchestrate regional immunity, we hypothesized that the therapeutic impact of CTLs originally sequestered around the periphery would differ from nascent or αPD1-activated CTLs in the systemic circulation. To this end, we examined the recruitment and trafficking of antigen-specific CTLs to tumors after vascular normalization and evaluated the proportion of systemic CTLs that reached tumors compared to pre-existing CTLs. We implanted GL261-MGH-SIINFEKL in CD45.1 mice and treated size-matched tumors with four i.v. doses of 1) control: IgG2a (250 µg/dose) + MOPC-21 (125 µg/dose), 2) αPD1 (250µg/dose), 3) A2V (125 µg/dose), and 4) αPD1+A2V (250 µg/dose+125 µg/dose) bi-weekly. Then we transferred 1 million OT-I cells to mice after the third dose (Fig. 4g). This timeline enables the expansion of OT-I T cells that can recognize OVA as a model tumor-associated antigen. The analysis of the TME in the excised tumors reproduced the vascular normalization (Fig. 1i-q), CTL re-localization (Fig. 4b), and CTL numbers and function measured in WT mice (Fig. 4d-f). Assessment of intratumoral CTLs revealed that endogenous CTLs (CD45.1) and OT-I cells (CD45.2) were significantly increased in tumors after vascular normalization with A2V alone or a combination of αPD1+A2V (Fig. 4h-i), consistent with the functional role of vascular normalization in effector CTL trafficking. Effector function through the production of IFNγ was largely dependent on αPD1 and was increased after αPD1+A2V therapy for both endogenous CD8 T cells and transferred OT-I cells (Fig. 4j-k). Collectively, these data (Fig. 2, Supplementary Fig. 7, and Fig. 4b-e) show that the contribution of CTL enrichment to antitumor immunity in CTL-excluded tumors depends on both functional vessels and reactivated CTLs with an ability to kill cancer cells.

Beyond T cell infiltration and activation, we assessed endogenous NK cell infiltration in GL261 and CT2A tumors (Supplementary Fig. 11). In GL261, A2V treatment increased intratumoral NK cells, with a further enhancement observed upon combination with αPD1. By contrast, neither treatment altered NK cell infiltration in CT2A.

Myeloid cells, specifically tumor-associated macrophages, constitute majority of the immune cell population in GBM ^24,25^. We and others previously reported that A2V reprograms tumor-associated macrophages from pro-tumor (M2-like, anti-inflammatory) to anti-tumor (M1-like, pro-inflammatory) in preclinical models of GBM^24,25^ and extra-cranial tumors ^47^. αPD1 monotherapy does not contribute to the polarization of macrophages from M2 to M1, but the effect of A2V on macrophage polarization was preserved in αPD1+A2V combination therapy. Since we observed cooperative interaction between myeloid cells and ECs in presenting antigen to CD8 T cells and the positive feedback loop through IFNγ production, we asked whether phenotypical changes in myeloid cells after therapy may contribute to IFNγ production. In concert with previous studies showing a broad spectrum of myeloid cells ^48^, we found that categorization of M1 versus M2 using classical markers may not be sufficient. We found that myeloid-derived suppressor cells (MDSCs, marked by GR1) produced cytokines that are not known for their suppressive function (*e.g.,* IL-12 and IFNγ; Supplementary Fig. 12). For these reasons, we analyzed the CD11b progenitor cells and defined M1-like cells as producers of inflammatory cytokines (such as IL-12 and IFNγ), and M2-like cells as producers of anti-inflammatory cytokines (such as IL-4 and IL-10) ^49^. αPD1+A2V therapy led to decreased expression of IL-10 and increased IL-12 expression (Supplementary Fig. 13). IL-12 plays a key role in linking innate and adaptive immune responses and is produced by myeloid cells functioning as antigen-presenting cells to instigate T cells to produce IFNγ ^50^. This interaction results in a positive feedback loop in favor of restoring antitumor immunity. We therefore asked whether IL-12 produced by myeloid cells after αPD1+A2V therapy directly contributes to IFNγ production by CTLs and also antigen-presentation by ECs (Supplementary Fig. 14). Blockade of IL-12 in a co-culture of CD8 T cells and ECs did not reduce antigen presentation by ECs or IFNγ production by CD8 T cells. However, in a co-culture of ECs, CD8 T, and myeloid cells, IL-12 blockade decreased EC’s ability to present antigen as well as IFNγ production by CD8 T cells. Addition of IL-12 to the co-culture of myeloid cells within first 24h after blockade rescued the phenotype (Supplementary Fig. 14, and Fig. 4l-o). Collectively, these data demonstrate that enhanced anti-tumor immune response after αPD1+A2V therapy can be attributed to the cooperative interaction between ECs and myeloid cells in presenting antigen to CTLs, which is partly mediated by IL-12.

### T-bet dependent durable response by CD8 T cells after αPD1+A2V therapy

Treatment with αPD1+A2V led to complete response in ∼20% of GL261-MGH and 005GSC bearing mice (Fig. 1i, k). To test the formation of immune memory in the tumor-free mice, we rechallenged these mice with GBM tumors in the contralateral hemisphere ^15^ (Fig. 5a). Indeed, GL261-MGH complete responders rejected contralateral GL261-MGH tumors (Fig. 5b), evidenced by the lack of visible and/or microscopic tumors 60 days after re-challenge. 005GSC complete responders developed contralateral 005GSC tumors with significantly slower kinetics when compared with 005GSC naïve mice (Fig. 5c). Since memory CD8 T cells are mediators of long-lasting immunity ^51^, we tested if there was a systemic CD8 T memory formation in the cured GL261-MGH mice (Fig. 5d). We found that CD8 T cells in cervical lymph nodes of cured mice had acquired a functional phenotype (CD44+) and produced increased effector cytokines (IFNγ + TNFα+) (Fig. 5e). Additionally, we observed an enhanced presence of central memory CD8 T cells (TCM) CD44+CD62L+CD127+ in two GBM tumor models (GL261-MGH tumors and CT2a tumors), and effector memory (TEM) CD44+CD62L-cells in the CT2a model after αPD1+A2V therapy in (Supplementary Fig. 15). Importantly, the increased accumulation of functional CD8 T cells was observed in the cervical lymph node (Fig. 5e) but not in the spleen (Fig. 5f), suggesting the formation of a local antitumor memory response.

**Figure 5.**
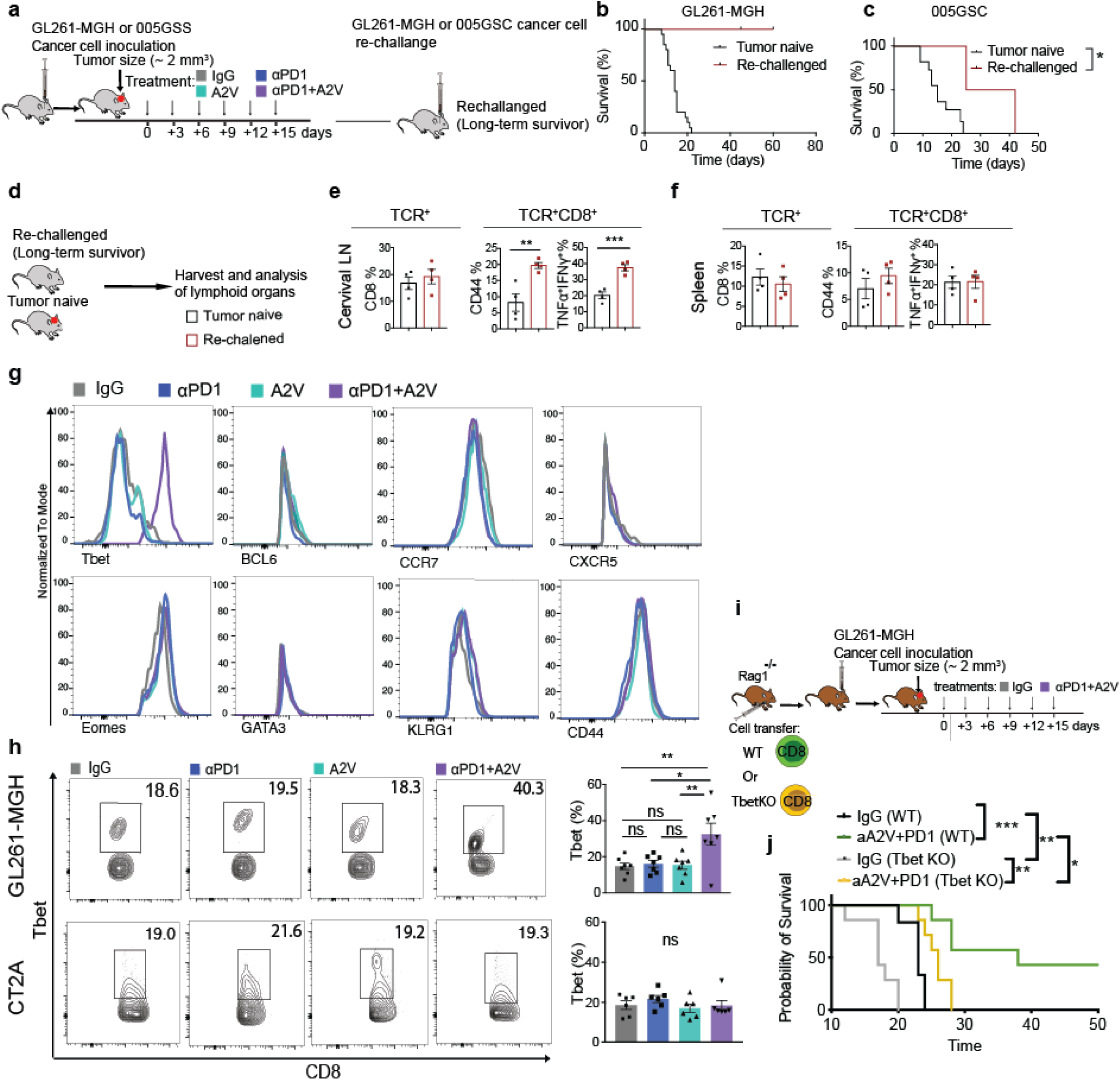
Induction of T-bet after αPD1+A2V regulates memory T cell formation in GBM. **a**) Schematic representation of the rechallenge experiment in the long-term survivors. **b-c**) Kaplan-Meier survival curves. αPD1+A2V generates a durable antitumor response in CTL-excluded tumor model GL261-MGH (n=4) and delayed tumor growth in 005GSC (n=4). **d-f**) Analysis of CD8 T cell numbers and phenotype in responders to αPD1+A2V treatment during memory response. CD8 T cell number and surface phenotype and cytokine production in cervical LN (e) and spleen (f) from GL261-MGH experienced mice versus naïve B6 mice shown after rechallenge with GL261-MGH. n=4. *\*P*<0.05. **g)** Histograms showing the expression of markers associated with CD8 T cell memory in GL261-MGH tumors (n=6). **h)** Assessment of the expression of T-bet in GL261-MGH (n=7) and CT2A (n=6). CTLs in GL261-MGH with a superior response to αPD1+A2V express significantly higher T-bet compared with monotherapies. GL261-MGH *P*=0.031 and CT2A *P*=0.4993. Multiple comparisons using the Tukey adjustment. For between group analysis post-Tukey, Statistical significance is shown as \**p* < 0.05, \*\**p* < 0.01, \*\*\**p* < 0.001, \*\*\*\**p* < 0.0001. **i)** Transfer of CD8 T cells (1 million) from WT or *T-bet* KO mice to *Rag1^-/-^* mice followed by therapy, as shown above. (**j**) Kaplan-Meier survival curves. Survival of mice after GL261-MGH inoculation is compared between WT CD8 or *T-bet* KO CD8 transferred to Rag1-/-hosts after IgG or αPD1+A2V treatments. Median survival of mice that received WT CD8 T cells was 23 days for IgG treatment (n=9) and 38 days for αPD1+A2V (n=7), Median survival of mice that received t *T-bet* KO CD8 T cells was 17 days for IgG treatment (n=7) and 25.5 days for αPD1+A2V (n=8). The survival curve compared using the Mantel cox test, *P*=0.0001 (WT) and *P*=0.003 (*T-bet* KO).

We investigated whether the difference in therapeutic outcome of GL261-MGH tumors compared with other GBM models could be attributed to the CD8 T cell response. Since there was a significant difference in survival outcome after αPD1+A2V therapy (Fig. 1i-j), we chose GL261-MGH and CT2A tumor models to address this question. Analysis of the TME showed CD8 T cells in GL261-MGH tumors produced low-to-medium levels of IFNγ +TNFα+, whereas CD8 T cells in CT2A produced high levels of Gzmb (Fig. 4f). Given that GzmB production marks terminally differentiated effector CD8 T cells, as opposed to IFNγ producing cells ^52^, the differences in the ability of CD8 T cells to convert into a memory phenotype in GL261-MGH versus CT2A may explain the differential response of these tumors to the same treatment.

We next examined specific markers associated with CD8 T effector and memory phenotype (T-bet, BCL6, CCR7, CXCR5, Eomes, GATA3, KLRG1, and CD44) (Fig. 5g). Among these, T-bet was prominently upregulated in CD8 T cells from GL261-MGH tumors that received αPD1+A2V therapy but remained unchanged in CD8 T cells from CT2A tumors (Fig. 5h). To test whether T-bet was required for durable CD8 T cell-mediated protection against GBM tumors, we transferred CD8 T cells from WT or *T-bet* KO mice to *Rag 1^-/-^* hosts followed by implantation of GL261-MGH cancer cells. Mice were then randomized and treated with control (IgG) or αPD1+A2V (Fig. 5i). GL261-MGH tumor bearing *Rag1^-/^*^-^ mice that received *T-bet*-WT but not *T-bet* KO CD8 T cells responded to αPD1+A2V therapy (Fig. 5j), indicating that T-bet is required for the generation of durable protective CD8 antitumor immunity consistent with a memory immune response.

## Discussion

GBM vasculature is often hyperpermeable and poorly perfused, resulting in vasogenic edema, hypoxia, low pH and a highly immunosuppressive tumor microenvironment (TME) ^2,8,10,12,24,25^. Thus, abnormal GBM vessels not only generate both physical and functional barriers to trafficking and extravasation of nascent and effector CD8 T cells, but also impair their function after they accrue in the TME, potentially contributing to the failure of all randomized phase III trials of ICBs in GBM patients ^1,53–57^. Here, we propose that vascular normalization using A2V can overcome these barriers by improving EC integrity resulting in improved blood flow required for delivery of cells to tumor vessels and inducing expression of ECAMs on GBM vessels required for CD8 T cell engagement and extravasation. Beyond increased trafficking of CTLs, we also show that combined αPD1+A2V can reprogram the GBM ECs into quasi-antigen presenting cells in both murine and human model systems and provide the underlying mechanism. Finally, we show that this approach is likely to have reduced immune related adverse events (irAEs) due to immune memory localized to tumor (Fig. 6).

**Figure 6.**
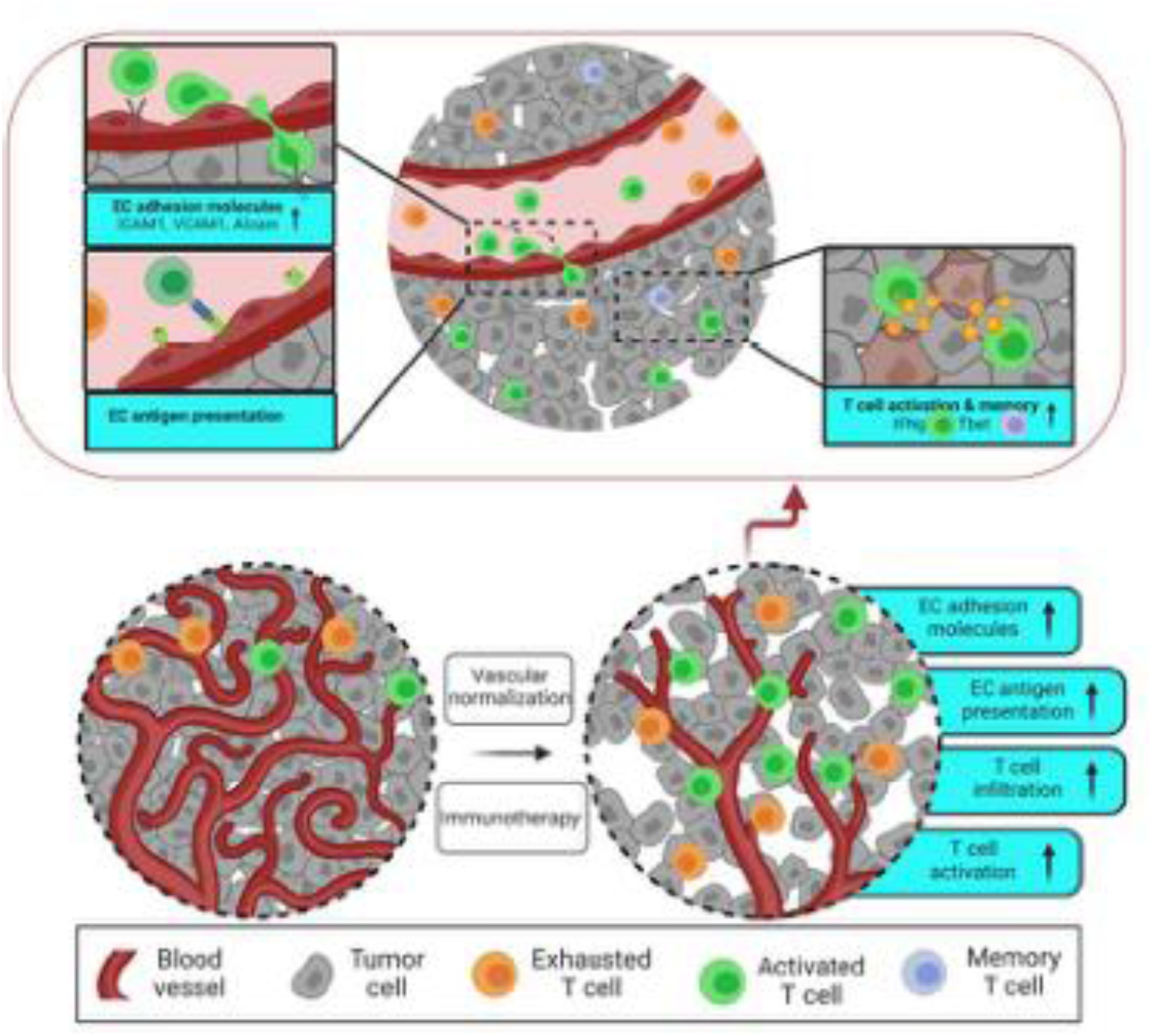
Mechanisms of benefit from concurrent blockade of Ang2, VEGF, and PD1 in GBM. We show here that GBM resistance to anti-PD1 and anti-VEGF therapy can be alleviated by concurrent blockade of Ang2. Combined blockade of Ang2, VEGF, and PD1 upregulates receptors required for effector CD8 T cells to infiltrate GBM and enables tumor endothelial cells to present tumor antigen to CD8 T cells, potentially reducing egress. The resulting durable anti-tumor response to combination therapy requires upregulation of T-bet in CD8 T cells. Finally, Memory CD8 T cells are sequestered in the draining lymph nodes after treatment, reducing the potential for immune-related adverse events.

A recent study ^58^ reporting therapeutic modality achieving vessel normalization showed transcriptomic changes in ECs that are distinct from our findings. The differences in genetic signatures of ECs treated with αPD1+A2V versus αPD1+aflibercept+AMG386 are likely due to different potency and mechanisms of action of the therapeutic compounds and the tumor models used. Unlike A2V that blocks both VEGF and Ang2, aflibercept traps VEGF-A, VEGF-B and PlGF, and AMG386 blocks both Ang1 and Ang2. Since Ang1 counters Ang2-signaling, combination of αPD1+aflibercept+AMG386 likely did not result in a durable anti-tumor response, unlike our study. Additionally, our experiments were performed with a GL261 derivative (GL261-MGH) that is resistant to αPD1 therapy, recapitulating the clinical scenario, compared to GL261 which is responsive to αPD1 therapy. This may also contribute to the different outcomes in gene expression and durable response in the two studies ^58^.

High endothelial venule (HEV) formation marked by presence of peripheral lymph node addressin (PNAd) on ECs in tumors has been shown to enhance infiltration and function of CTLs ^59^, predict immunotherapy response ^60^, and aid response to immunotherapy in extracranial tumors ^61^. In contrast to these studies ^60,62^, immunostaining of ECs in our GBM models did not show significant level of PNAd expression (Supplementary Fig. 10a). Using flow cytometry, we also found negligible frequency of ECs expressing PNAd (Supplementary Fig. 10c). This discrepancy may be due to specific features of GBM cells and/or subcutaneous tumor model used in other studies ^59,60^, as the subcutaneously grown tumors (ectopic) are known to be different from their orthotopic counterparts ^63^. Moreover, PNAd plays a critical role in initial tethering and rolling of naive T cells for trafficking through the lymphatic system ^64,65^. However, T cells infiltrating into the tumors include effector and/or memory cells that may need different adhesion molecules. In fact, we found upregulation of *ALCAM* at mRNA and protein levels after αPD1+A2V therapy, which may support effector T cell trafficking into the tumor ^66^ (Supplementary Fig. 10g-h). Collectively, these data suggest αPD1+A2V therapy reprograms GBM ECs to express appropriate ECAMs to enable trafficking of CTLs into GBM.

In the process of T cell migration into tumor, ECs may interact with T lymphocytes in an MHC-dependent fashion. Professional antigen presenting cells, such as dendritic cells, exert their canonical function by (1) constitutive MHC class I and MHC class II expression in combination with the ability to efficiently ingest and process antigens; (2) provision of proper co-stimulation to prime naïve CD4 or CD8 T lymphocytes; and (3) induction and promotion of T lymphocyte differentiation and function. Microvascular ECs in organs, such as, liver and heart appear to fulfill these criteria ^67^. These ECs may function as semi-professional antigen presenting cells, because they do not express certain co-stimulatory molecules, yet they clearly have the capacity to induce T cell responses *in vitro* and *in vivo.* While most of these responses are known to induce T cell anergy, ECs can induce inflammatory responses in certain immunological conditions such as malaria infection ^68^. We discovered here that blockade of Ang2, VEGF, and PD1 reprogramed ECs to present tumor antigen to CD8 T cells in GBM and consequently activated T cells that show anti-tumor activity. A2V alone was able to induce IFNγ response-related genes in blood vessels (Fig. 2c-e, Supplementary Fig. 5). However, the combination (αPD1+A2V) therapy was required for cross presentation of antigen to CD8 T cells in GBM (Fig. 2), similar to the ECs in malaria infection ^69^. Moreover, we found a positive feedback loop between IFNγ produced by CTLs and elevated IFNRs on ECs after αPD1+A2V therapy may lead to enhanced antigen presentation and T cell activation. This is in line with the recent finding of IFNγ responsiveness of host immune cells ^70^ and IFNγR signaling in solid tumors, specifically in GBM, enabling antitumor function of endogenous CTLs and exogenous CAR-T cells ^46^. This positive feedback loop may be important because the phenotype of ECs may be sustained by the local organ-specific microenvironment. To maintain the EC barrier during inflammatory conditions, ECs may need to remain constitutively resistant to cell death. Indeed, a key feature of ECs is the resistance to inflammatory stimuli activating the extrinsic pathway of apoptosis ^71–73^. Akt activation by CD31 signals prevents the translocation of the forkhead transcription factor FoxO3 to the nucleus, thus inhibiting transcription of the proapoptotic genes CD95/Fas and caspase 7, and derepressing the expression of the antiapoptotic gene cFlar ^74^. This feature lowers the risk of CTL-mediated immune attack against ECs.

In view of the ability of normalized ECs to present antigens as well as promote trafficking of effector and memory CTLs into the tumors, the egress of CTLs may be a driver of limited therapeutic outcome. The significantly higher therapeutic outcome observed in GL261-MGH compared to CT2A tumors after αPD1+A2V therapy may be explained by distinct localization and effector cytokine phenotype of CD8 T cells. CT2A tumors are CTL-included, and the CTLs enhance their effector function after therapy, evidenced by high levels of GzmB and IFNγ. CTLs in GL261-MGH are in the periphery and localize into tumors only after therapy with A2V or αPD1+A2V. CTLs that enter GL261-MGH after αPD1+A2V produce significantly lower levels of GzmB compared with CT2A (Fig.4f). These data indicate that the differentiation stage of CTLs is likely different in these tumors. While GL261-MGH CTLs may be in early differentiation stage, CT2A CTLs likely reached the terminal differentiation stage and may not have the capacity to convert into the effector/memory subtype, and thus a durable response could not be realized.

The observation that a localized memory response restricted to dLNs was generated after αPD1+A2V therapy that does not extend systemically, suggests that this drug combination may have less irAEs - a significant challenge in immunotherapy of cancer ^75^. irAEs often lead to loss of therapeutic benefit due to treatment discontinuation ^76^. Moreover, therapeutic interventions to control irAEs include administration of corticosteroids to control inflammatory and vasogenic edema ^13,35,76^. Because A2V controls edema without affecting αPD1 mediated inflammatory responses, the combination therapy αPD1+A2V may be a rapidly translatable option due to reduce irAEs in GBM patients.

The combination of αPD1 with A2V has several advantages over conventional methods that rely on vascular normalization by αVEGF alone or by ICB therapy alone ^2^. First, acquisition of antigen presentation capacity by ECs can promote activation of CD8 T cells. Second, αPD1 can maintain effector T cell status including IFNγ production, which may result in feedforward response towards CTL trafficking to tumors. Third, targeting resistance mechanisms to anti-VEGF/VEGFR therapies with A2V may extend the duration of vascular normalization, thereby facilitating the enrichment of functionally competent CTLs within GBM and enhancing their antitumor activity, with the potential to promote the formation of a durable memory T cell pool.

Collectively, our data (i) provide heretofore unknown insights into targeting Ang2 as a shared resistance pathway for both αVEGF and αPD1 in experimental GBM models, (ii) demonstrate a strategy to reprogram ECs to activate and promote CTL recruitment into GBMs followed by retaining intratumoral T cells, (iii) offer a possible solution to overcome GBM resistance to αVEGF and αPD1, and (iv) provide mechanistic insights into how T cell memory forms after αPD1+A2V therapy (Fig. 6). Multiple strategies to combine αPD1 and αVEGF are currently being evaluated in clinical trials for non-CNS tumors and 7 such combinations have been approved by the US-FDA for lung, kidney, liver and endometrial cancers ^2^. Our study provides a foundation for testing the combination of a more effective vascular normalizer A2V in combination with αPD1 in GBM patients where αPD/L-1 monotherapy and in combination with αVEGF have failed in all randomized clinical trials. A2V may also improve the outcome of CAR-T cells beyond anti-VEGF for GBM ^20^.

## Methods

### Antibodies, chemicals, peptides, recombinant proteins, and assay reagents

For ease of identification, full details of antibodies and materials are shown under the registry of publicly available database (antibodyregistry.org) that provides a comprehensive list of suppliers, and appropriate references for usage. This information is summarized under Identifier in Supplementary Table 1.

## Supplementary Methods

### Antibodies, chemicals, peptides, recombinant proteins, and assay reagents

**Supplementary Table 1:**
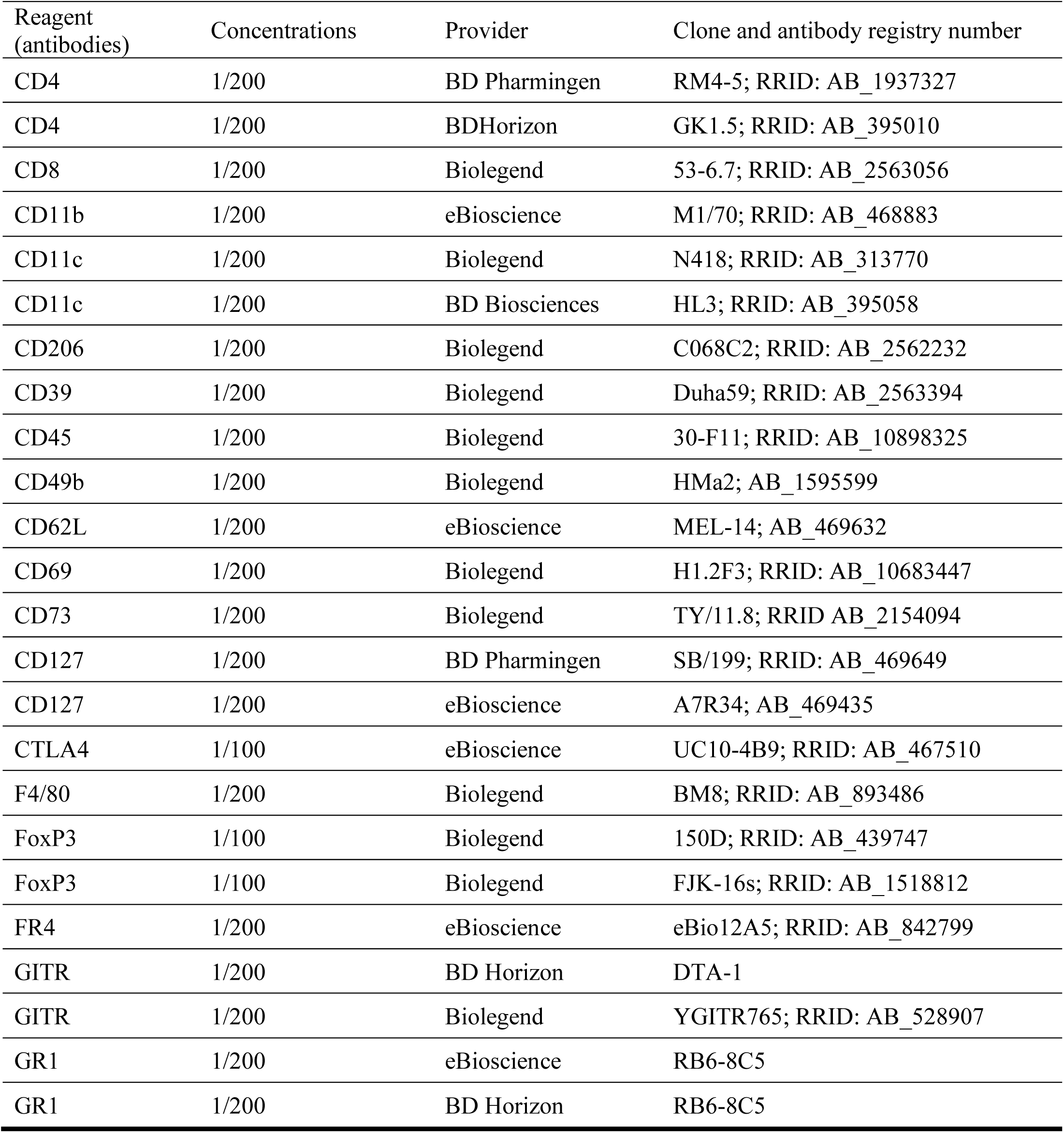
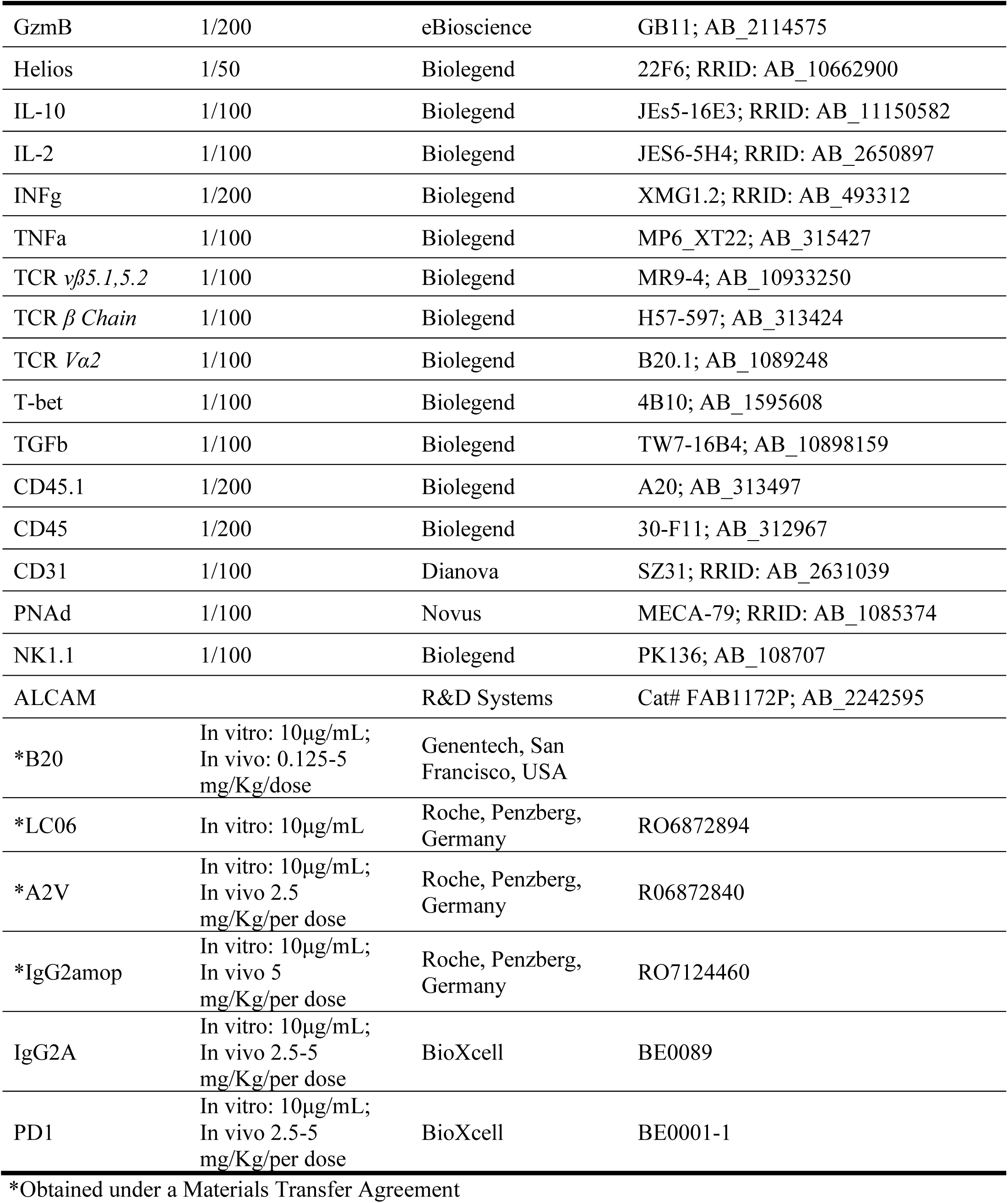
Antibodies List. For ease of identification, full detail of antibodies and materials are shown under the registry of publicly available data base (antibodyregistry.org) that provides a comprehensive list of suppliers, and appropriate references for usage. This information is summarized under antibody registry number in the listed table.

#### Cell Culture

The murine GL261-MGH GBM cell line ^15^ was grown in serum-free conditions using the NeuroCult NS-A proliferation kit (Stemcell Technologies). The murine CT2A was obtained from Dr. Thomas N. Seyfried’s laboratory at Boston College and cultured in Dulbecco’s Modified Eagle Medium (DMEM) supplemented with 10% fetal bovine serum and 1% penicillin-streptomycin (10,000 U/mL). 005GSC cells were obtained from Dr. Samuel D. Rabkin’s laboratory at MGH. The GL261-MGH and CT2A cell lines expressing secreted Gaussia luciferase (GLUC) were generated by transducing cells with a lentiviral vector co-expressing GLUC and GFP, provided by the MGH Vector Core, followed by sorting. The 005GSC cell line expressing secreted GLUC was generated by transfecting cells with a GLUC plasmid followed by puromycin selection. All cell lines were grown in a humidified atmosphere of 5% CO2 and 95% air at 37°C and repeatedly tested and found negative for mycoplasma using the Mycoalert Plus Mycoplasma Detection Kit (Lonza).

#### SIINFEKL expressing cell lines

Lentiviral vectors encoding SIINFEKL were a kind gift from Dr. Arlene Sharpe (Harvard Medical School). Lentiviral particles were packed by co-transfecting transfer plasmid with pSPAX2 pMD2.G into 293T cells. Supernatant was harvested and incubated with GL261-MGH cells in the presence of polybrene. Transduced cells that stably express SIINFEKL were selected using puromycin (5ug/ml).

#### Flow Cytometry

Single-cell suspensions were prepared from each organ [tumor, lymph node (LN), spleen, and blood] of tumor-bearing mice. Specifically, tumor-only regions were isolated under the stereotactic microscope followed by single cell preparation. Cells were stained with antibodies listed in the Core Resource Table (Suppl Table 1) with concentrations of 1μg/ml for all targets. Surface staining was performed on ice for 20 min. For cytokine expression analysis, cells were activated with Leukocyte Activation Cocktail (BD Biosciences, Cat no: 554656) in RPMI containing 10% FBS and 1% penicillin-streptomycin (10,000 U/mL) for 4.5 hr. Cells were first stained with surface markers in FACS buffer (2% BSA in PBS) for 20 min on ice. Then cells were fixed in Fix/Perm buffer (eBioscience) for 15 min, washed in permeabilization buffer (eBioscience), and stained for intracellular factors in permeabilization buffer for 20 min on ice. Live cells for transfer experiments and *in vitro* assays were sorted using BD FACSAria™ II cell sorter (BD Biosciences). Cells were analyzed on Aria II (BD Biosciences) or LSRII (BD Biosciences), and data analysis was performed on FlowJo (Tree Star 10.7v).

#### Tumor Models

C57BL/6, *OT-1*, *Rag1^-/-^*, and *T-bet* knockout mice were obtained from the Jackson laboratory and maintained in the Cox-7 gnotobiotic animal facility operated by the Edwin L. Steele Laboratories, Department of Radiation Oncology at the MGH. All animal experiments were performed in the Cox-7 animal facility, accredited by the Association for Assessment and Accreditation of Laboratory Animal Care International (AAALAC). Female and male mice were used. Animal protocols were approved by the Institutional Animal Care and Use Committees (IACUC) at MGH. C57BL/6 or *Rag1^-/-^*mice were injected with cancer cells (1×10^5^ GL261-MGH-GFP-GLUC, 5×10^4^ CT2A-GFP-GLUC, 7.5×10^4^ 005GSC-GFP-GLUC) orthotopically using a stereotactic device in the forebrain. Tumor size was measured either by micro-ultrasound, blood GLUC, or both. To ensure GFP-Gluc cancer cells are non-immunogenic, we previously compared their growth with parental lines (Method Figure 1) ^15^. Tumors were size-matched and randomized to treatment groups. After treatment initiation, tumor size was monitored every 3 days by blood GLUC measurements.

**Method Figure 1.**
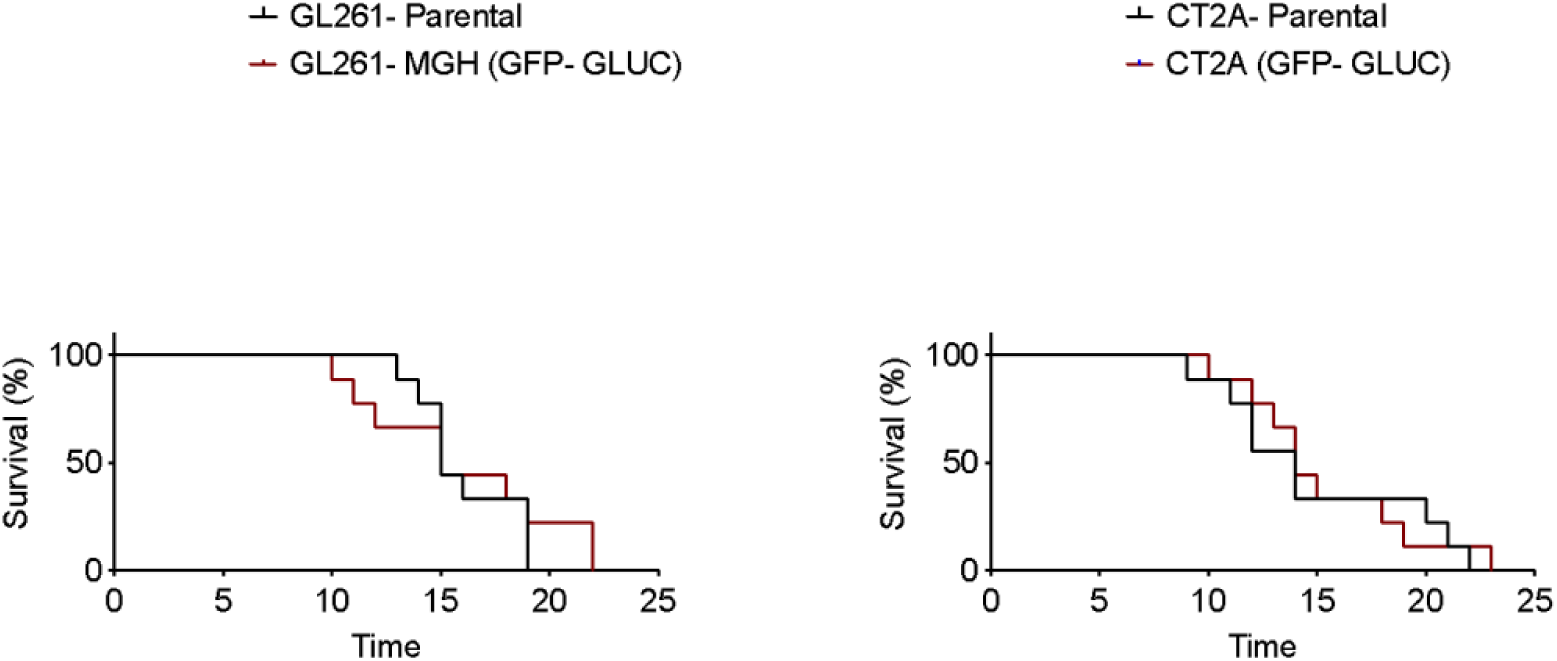
Average survival days after tumor randomization for parental GL261 (n =9, ∼15 days) and CT2A tumors (n=9, ∼14 days) were similar to GFP-Gluc version of each after randomization.

For survival studies, C57BL/6 mice bearing GL261-MGH-GFP-GLUC, CT2A-GFP-GLUC, or 005GSC-GFP-GLUC were treated intraperitoneally (i.p) every 3 days with total 6 doses of i) IgG2a; ii) αPD1 (Bioxcell, RMP1-14); iii) B20 (Genentech); or iv) αPD1+B20. Another set of mice was treated every 3 days with total 6 doses of i) IgG2a; ii) αPD1 (Bioxcell, RMP1-14); iii) A2V (Roche); or iv) αPD1+A2V. Tumor size was measured every 3 days until the first mouse showed clinical signs of GBM associated morbidity (including serious movement problems, hunch-back, and/or weight loss beyond 15%), at which time the mice were euthanized.

C57BL/6 mice bearing GL261-MGH-GFP-GLUC were treated every 3 days with total 4 doses of i) IgG2a; ii) B20 1.25 mg/kg body weight; iii) B20 2.5 mg/kg body weight; or iv) B20 5 mg/kg body weight. The mice were also injected with 60 mg/kg of pimonidazole at 10 mg/ml 1 h before tumor removal for hypoxia studies. For vessel perfusion analysis, mice were slowly injected with 100 ml of 1 mg/ml biotinylated lectin (Vector Labs) via the retro-orbital sinus 5 min before tumor removal. The tumors were then excised, fixed in 4% formaldehyde in PBS, followed by incubation in 30% sucrose in PBS overnight at 4 C and frozen in optimal cutting temperature compound (Tissue-Tek). Transverse tumor sections, 40 mm thick, were immunostained with antibodies to endothelial marker CD31 and counterstained with 40,6-diamidino-2-phenylindole (Vector Labs).

*Rag1^-/-^* mice received i) vehicle, ii) WT 1×10^6^ CD8 T cells (TCR^+^CD8^+^), iii), or iv) *T-bet* KO 1×10^6^ CD8 T cells (TCR^+^CD8^+^). Mice were inoculated with GL261-MGH-GFP-GLUC two days after cell transfer. For each set of cell transfer experiments, mice were randomized to two cohorts post tumor establishment (size 0.5-1 mm^3^) for treatment with IgG2a or αPD1+A2V (n=6-9 per cohorts). They were then treated with 6 doses of each antibody (250 µg/mouse/dose). Tumor size based on blood GLUC was measured until the first mouse showed clinical signs of GBM associated morbidity, as stated above.

For immunological assays, mice were sacrificed 2-3 days after the fourth treatment. Tumor, LNs, blood and spleen were harvested. Single-cell preparation (including TILs) was conducted using a modified mechanical dissociation method, and cells were analyzed as previously described in the antibody and flow cytometry section.

For multi-arm treatment studies, 8-12 mice were assigned to each group to achieve statistical power. For two-arm cell transfer experiments, at least 4-5 mice were used in each group. Tumor growth was analyzed using two-way ANOVA with multiple comparisons. Event-free survival (moribund) estimates were calculated with the Kaplan-Meier method. Groups of mice were compared by log-rank test.

Brain edema was quantified by measuring tissue water content using the wet-to-dry weight method. Following euthanasia, brains were rapidly harvested and weighed to obtain the wet weight. Tissues were then dried at 100°C for 24–48 h until a constant weight was achieved and reweighed to obtain the dry weight. Brain water content was calculated as: [(wet weight − dry weight)/wet weight] × 100.

#### Tumor blood vessel RNA isolation and RNA-Seq Analysis

RNA was isolated from tumor blood vessels of treated mice. Briefly, tumor-only regions were isolated under the stereotactic microscope. Brain tissue into small pieces and using Dounce homogenizer on ice. Tumor blood vessels were separated using 15% Ficoll-solution by centrifugation. Pellets were resuspended in 1% BSA and put in a 40 μm cell strainer on top of a 50ml Falcon. Debris was washed through the strainer, and purified blood vessels were eluted from the strainers. Purity and the integrity of blood vessels were examined under microscopy. RNA was isolated using the RNeasy Mini Kit (Qiagen). The examination of RNA integrity, concentration, library preparation and sequencing were performed by NGS core at MIT.

#### Bioinformatic analysis

FastQC was used for the quality control of raw sequencing data. After quality control, Cutadapt was used to remove the low-quality bases and adaptor contaminations. The quality of yield clean data was examined by FastQC software again. Next, Hisat2 was used to align the clean data to mouse reference genome mm10 which we downloaded from Illumina iGenomes database. After data mapping, samtools was used to manipulate SAM files and BAM files. HTSeq-count was used to count the number of reads that were aligned to the gene features. The differentially expressed genes were identified by DESeq2 with adjusted p value cutoff < 0.05 (Benjamini-Hochberg method). PCA analysis was performed by R “prcomp” function with top 500 most variable genes. GSEA analysis was performed by GSEA with standard genesets from MSigDB database.

#### *Ex Vivo and In Vitro* Co-Culture Assays and human tumoroids

For Organotypic brain slice culture, 200 μm thick brain slices were obtained from postnatal day 17 to 20 mice using a Compresstome Vf-300 microtome (Precisionary Instruments, Compresstome, VF-300). Slice cultures were grown on Hydrophilic PTFE cell culture inserts (Millipore) with 50% MEM, 25% EBSS, 25% HS, Gentamycin and Glucose (Invitrogen). Five thousand GL261-Gluc cells were injected into the cortical layers 12-24hrs after slice preparation. CD8 or CD4 T cells, enriched from Spleen of C57BL6 mice. To assess tumor growth, medium was collected every two days, and Gluc activity was measured with a Promega Glomax 96 microplate luminometer (Fisher Scientific). Immune cell number and phenotype were assessed by flow cytometry. The number of slices required to demonstrate statistical significance in each group was estimated based on findings from previous experiments. No data were excluded from the analyses

In preclinical models, Endothelial cells (CD45^-^CD31^+^), myeloid cells (CD11b+), and CD8 T cells were isolated from GL261-MGH tumors. Cell sorting was done post enrichment using CD31 isolation or CD8 isolation kits. Pre-sorting enrichment assays in mice are done using commercial kits (stem cell technologies). CD8 T cells were negatively selected using Easysep ® CD8a T Cell Isolation Kit, mouse (Stemcell technologies, 19853). CD11b cells were isolated using Easysep ® CD11b positive selection kit (Stemcell technologies, 18970). Using CD31-PE antibody, CD31 cells were tagged and then enriched using Easysep ®PE-positive selection kit (Stemcell technologies, 17666).

Culture media was Roswell Park Memorial Institute medium (RPMI) supplemented with 10% FBS and 1% penicillin-streptomycin. Ratio of CD8 to Endothelial cells to CD11b cells to cancer cells were 1:1:1:1. Cell number, phenotype and cytokine production were measured using flow cytometry within the first 3 days. For experiments with long-term culture (>3 days), IL-2 (20 ng/mL) was added to the culture.

For the hydrogel-embedding co-culture of human GBM cells and HUVECs (referred to as in vitro human tumoroids), HLA expression on various human GBM cells were first characterized. U-87, U-251, U373, and SNB19 cells are homozygous at HLA-A2, which may provide enhanced recognition, and represent roughly half of the general U.S population. These GBM cells expressing Nuclight Red using Lentivirus were sorted and then maintained under selection pressure pre-experiments. Cancer cells were cultured with high-glucose Dulbecco’s Modified Eagle Medium (DMEM; GenDEPOT) containing 10% fetal bovine serum (premium, US origin; GenDEPOT) and 100 U ml^−1^penicillin and 100 μg ml^−1^ streptomycin in a humidified 5% CO_2_ incubator. HUVECs (Promocell) were cultured with Endothelial Cell Growth Medium-2 (Lonza). The culture medium was changed every 2 or 3 days. Prior to the hydrogel-embedding culture, we harvested each cell type by treating with a solution of 0.25% trypsin in EDTA (Gibco) for 3 min, collected the cells by centrifugation at 1,500 r.p.m. for 5 min and then dispersed the cells in each culture medium. The cell suspensions were mixed with the pre-gel solutions (10 mg ml^−1^) of either BdECM or collagen to a final concentration of 5 × 10^6^ cells ml^−1^ for both cell types. The aliquots of each cell-encapsulated pre-gel solution were transferred to the wells of a multi-well plate and placed in the incubator at 37 °C for 1 h. After gelation, the culture medium for the cancer or endothelial cells was added to the wells. Each culture medium was changed every 2 or 3 days.

#### Quantification and statistical analysis

Statistics were performed using Prism v8. One-way analysis of variance (ANOVA) with Tukey’s post hoc test was used as indicated in the figure legends. The Kaplan-Meier method was used for survival studies as indicated in the figure legends. Student’s t-test was used for two arm studies as indicated in the figure legends. N represents the number of mice used in the experiment, with the number of individual experiments listed in the legend. Graphs show individual or in case of survival studies combined experiments/samples. Results are presented as mean with or without error bars showing the standard error of the mean (SEM). Differences with p < 0.05 were considered statistically significant.

## Data availability statement

All data generated by this study are included in the manuscript, extended data, supplementary files or source data files. Previously published microarray and RNA-sequencing data that were re-analyzed here are available in Gene Expression Omnibus (GEO) under accession code (GSE212223). Source data are provided with paper. Materials are available under a material transfer agreement (contact person RKJ).

## Acknowledgments/Financial Support

We thank Dr. Timothy Padera for his helpful comments on our manuscript, and Sylvie Roberge (MGH Boston) for outstanding small animal surgery support. DF’s work is supported through NIH grants R01-CA208205 and R01NS118929. RKJ’s work is supported through NIH grants R35-CA197743, R01-CA208205, R01-CA259253, R01-CA269672, U01-CA224173, U01-CA224348 and U01-CA261842, and by the National Foundation for Cancer Research, Harvard Ludwig Cancer Center, Nile Albright Research Foundation, and Jane’s Trust Foundation. T.P.U. is supported by T32 Training Grant 5T32CA272386.

