## Supplementary data for "Combined blockade of VEGF, Angiopoietin-2, and PD1 reprograms glioblastoma endothelial cells into quasi-antigen-presenting cells"

Current affiliations:
a. AstraZeneca, 290 Binney Street, Cambridge, MA 02142
b. AstraZeneca, 1 Medimmune Way, Gaithersburg, MD 20878.
c. MGB Cancer Institute Peppas Center for Neuro-Oncology, 33 Fruit Street, Boston, MA 02144.
d. Trutino Biosciences, 10179 Huennekens St #101, San Diego, CA 92121.
e. Ability Biotherapeutics, Montreal, Quebec.
f. AgenusBio, 3 Forbes Road, Lexington, MA 02421, USA.
g. Genentech, 1 DNA Way, South San Francisco CA 94080, USA.

[†] These authors contributed equally

**Supplemental Figures:**

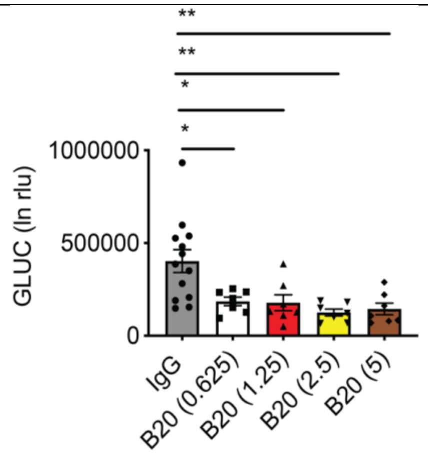


Supplemental Figure 1. Production of GLUC by cancer cells as a measure of cancer cell viability after therapy. GL261-MGH bearing mice was treated with IgG (n=13), B20 (0.625 mg/kg BW; n=7), B20 (1.25 mg/kg BW; n=7), B20 (2.5 mg/kg BW; n=7), and B20 (5 mg/kg BW; n=7). Data presented are mean ± SEM. $P$ =0.004 by ordinary one-way ANOVA test and were corrected for multiple comparisons using the Tukey adjustment. For between group analysis post-Tukey, Statistical significance is shown as $*p < 0.05$, $**p < 0.01$.

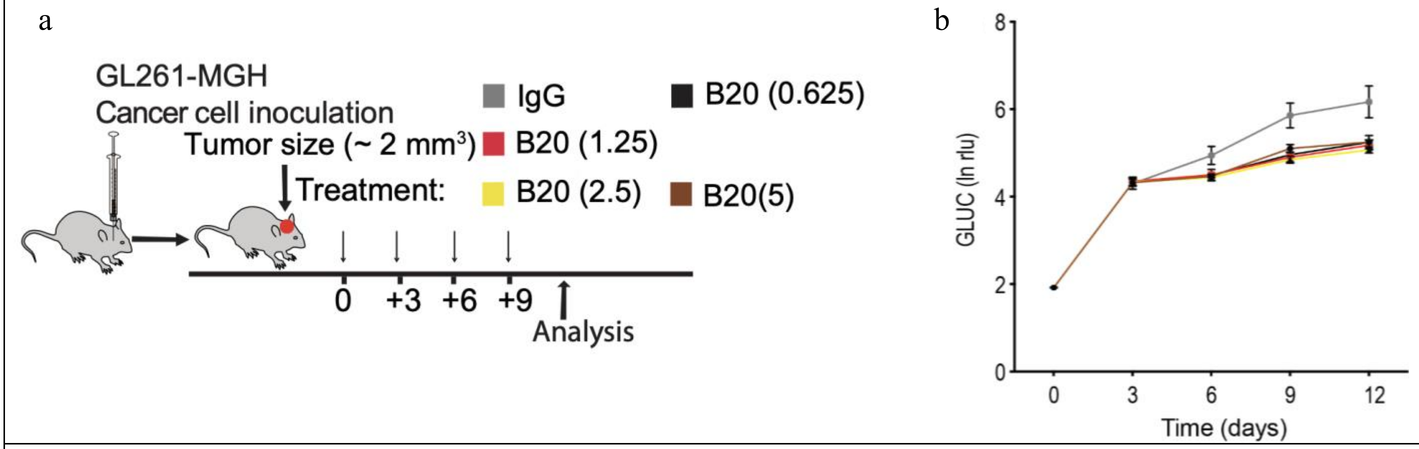


Supplemental Figure 2. GBM growth over time in response to different doses of B20. **a**) Schematic of treatment protocol. Mice were inoculated with $1x10^5$ GL261-MGH cells orthotopically and tumor growth was monitored using serial measurements of Gaussian luciferase activity (GLUC) in the blood. Mice were randomized into 4 treatment groups when tumor size reached ~2 $mm^3$ and then treated with IgG (n=13), B20 (0.625 mg/kg body weight (BW; n=7), B20 (1.25 mg/kg BW; n=7), B20 (2.5 mg/kg BW; n=7), and B20 (5 mg/kg BW; n=7). Doses were administered every 3 days intraperitoneally. **b**) Tumor growth curves were compared by two-way ANOVA (Days × Group). Multiple comparison showed significance versus IgG for all groups [$P$=0.0005 for IgG vs. B20 (0.0625); $P$=0.0005 for IgG vs. B20 (1.25); $P$=0.0004 for IgG vs. B20 (2.5); $P$=0.0005 for IgG vs. B20 (5)].

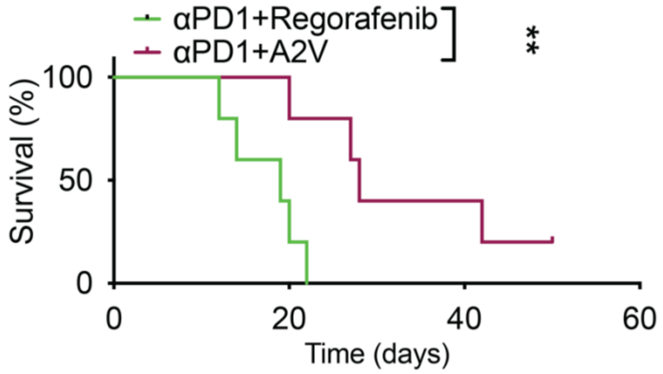


Supplemental Figure 3. Survival outcome of GL261-MGH bearing mice treated with αPD1+A2V (n=5) versus αPD1+ Regorafenib (n=5). Median survival tumor-bearing mice treated with αPD1+A2V (n=5) was 28 days versus 19 days for αPD1+ Regorafenib (n=5). Survival curve comparison done using Mantel Cox test, $P$= 0.0088.

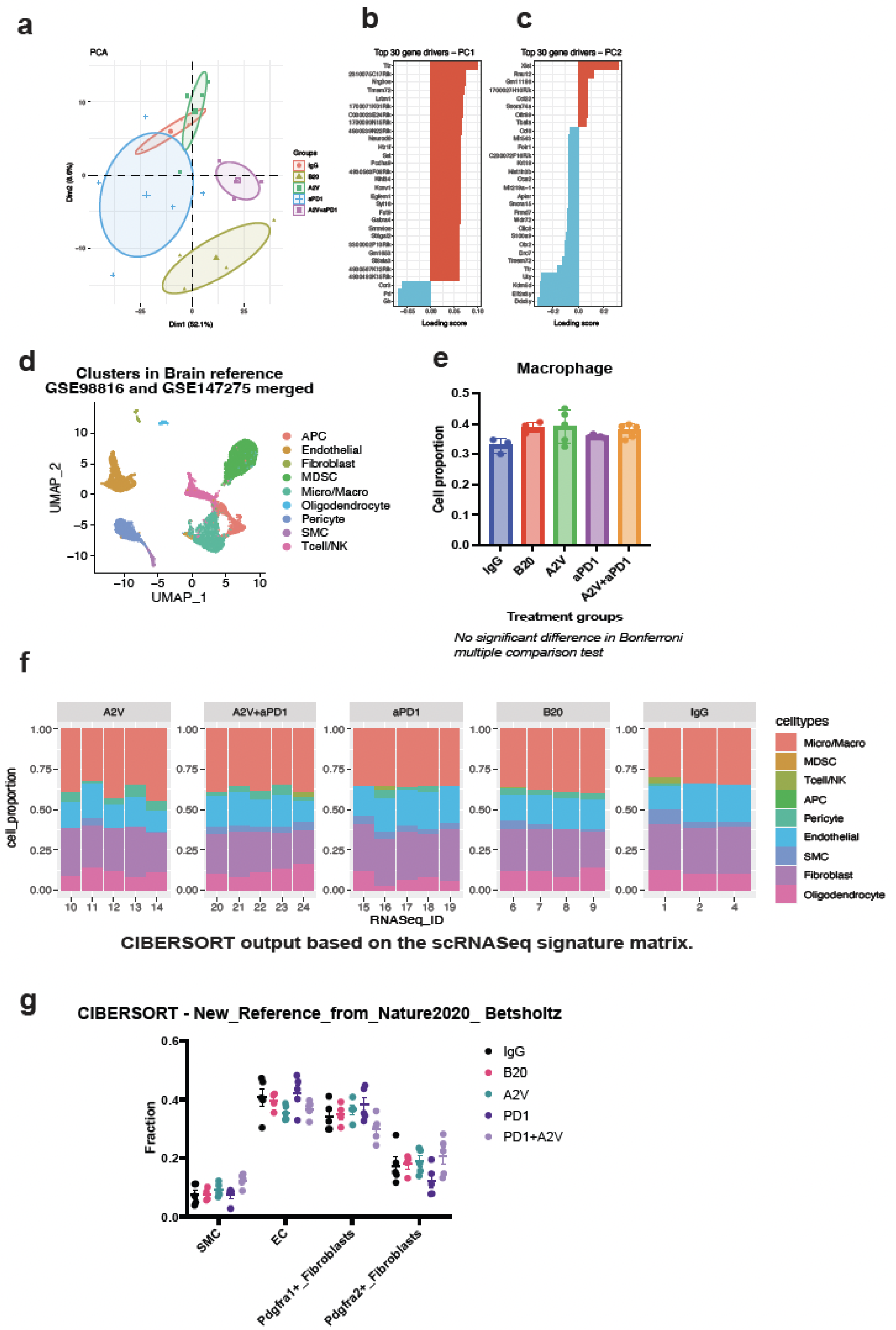
a
PCA
Groups
IgG
B20
A2V
aPD1
A2V+aPD1
Dim1 (52.1%)
Dim2 (8.6%)
b
Top 30 gene drivers – PC1
Loading score
c
Top 30 gene drivers – PC2
Loading score
d
Clusters in Brain reference
GSE98816 and GSE147275 merged
UMAP_1
UMAP_2
APC
Endothelial
Fibroblast
MDSC
Micro/Macro
Oligodendrocyte
Pericyte
SMC
Tcell/NK
e
Macrophage
Cell proportion
IgG
B20
A2V
aPD1
A2V+aPD1
Treatment groups
No significant difference in Bonferroni multiple comparison test
f
A2V
A2V+aPD1
aPD1
B20
IgG
cell_proportion
RNASeq_ID
celltypes
Micro/Macro
MDSC
Tcell/NK
APC
Pericyte
Endothelial
SMC
Fibroblast
Oligodendrocyte
CIBERSORT output based on the scRNASeq signature matrix.
g
CIBERSORT - New_Reference_from_Nature2020_ Betsholtz
Fraction
SMC
EC
Pdgfra1+_Fibroblasts
Pdgfra2+_Fibroblasts
IgG
B20
A2V
PD1
PD1+A2V

Supplemental Figure 4. CIBERSORT Analysis. (a) PCA plots of GBM treatment groups. (b-c) Top 30 gene drivers for PC1 and PC2 axis. (d-f) Clustering of 2 brain specific scRNAseq datasets (GSE98816 and GSE147275) to generate the reference gene signature for CIBERSORT analysis. (e) Comparison of macrophage-like features across treatment groups. (f) Cell proportions identified in bulk RNA sequencing samples. (g) The deconvolution of bulk RNA-Seq data with normal brain vasculature signatures.

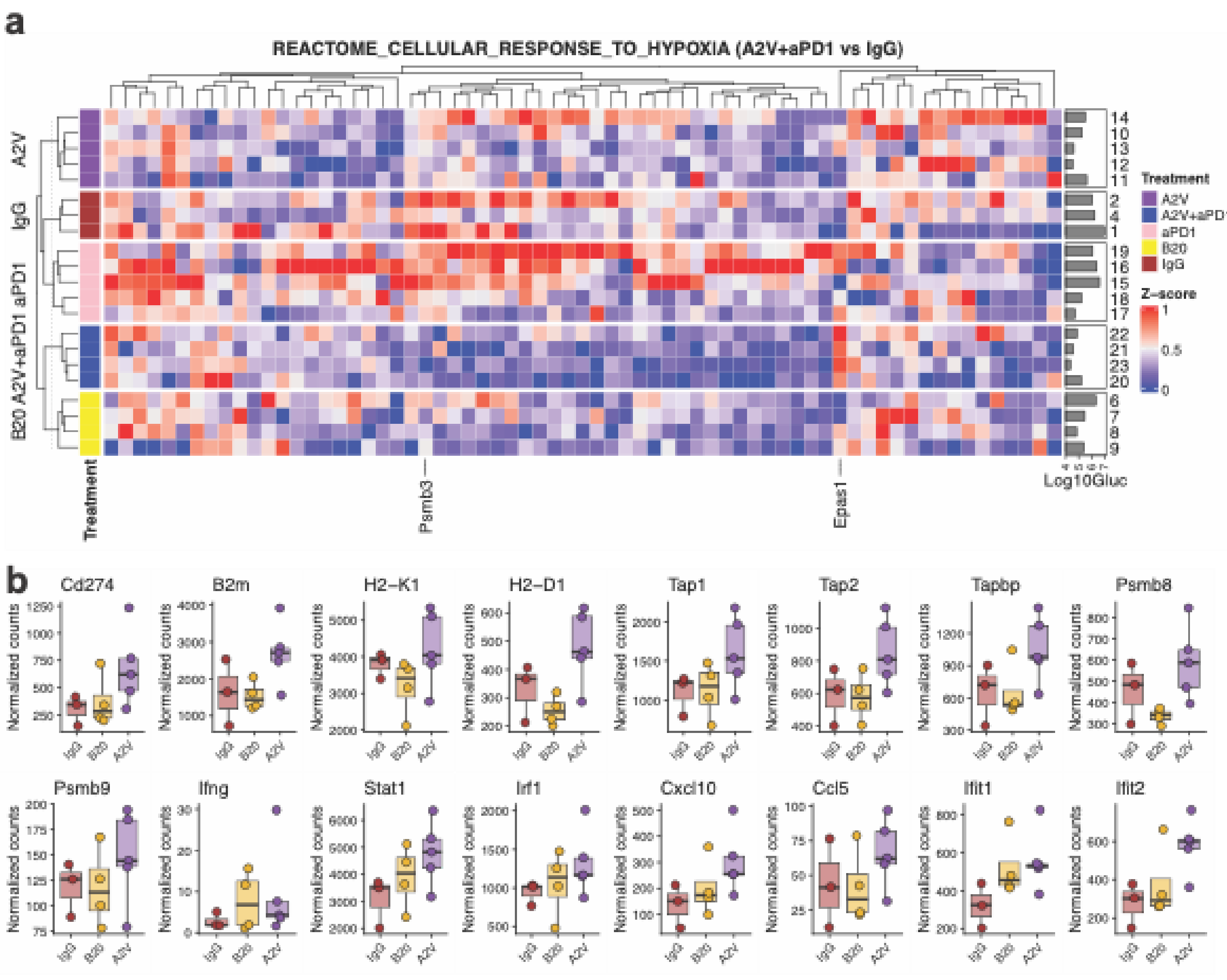


Supplemental Figure 5. (a) Heatmap of genes in the REACTOME_CELLULAR_RESPONSE_TO_HYPOXIA gene set, which was significantly enriched (reduced) in A2V+aPD1 tumors compared to IgG. (b) Normalized counts for individual interferon-γ pathway genes across IgG, B20, and A2V treatment groups.

| | |
|---|---|
| 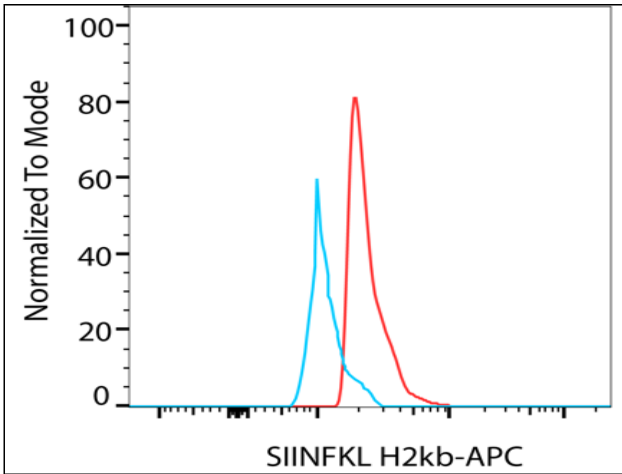 | Supplemental Figure 6. Flow cytometric measurement of expression of SIINFEKL on H2K$^b$ (mice MHC class I) on GL261-MGH (mouse). Blue: isotype control. Red: anti- SIINFEKL H2kb. Data is representative of 3 biological replicates. |

| | |
|---|---|
| 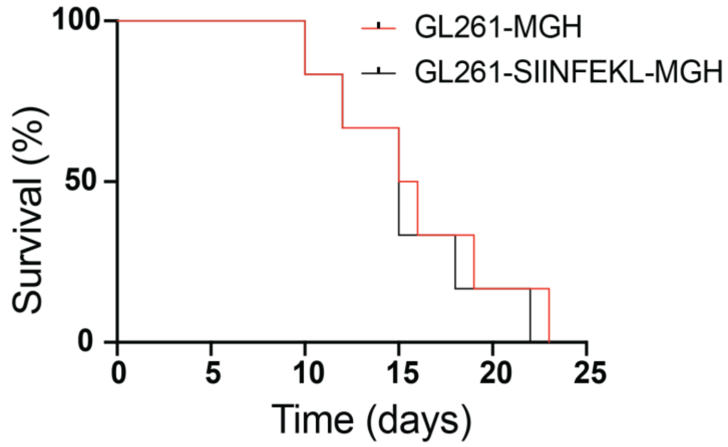 | Supplemental Figure 7. Median survival of GL261-MGH (n=6) was 15.5 days and GL261-SIINFEKL-MGHGL261-SIINFEKL-MGH (n=6) was 15 days. Survival curve comparison done using Mantel cox test showed no significant difference. |

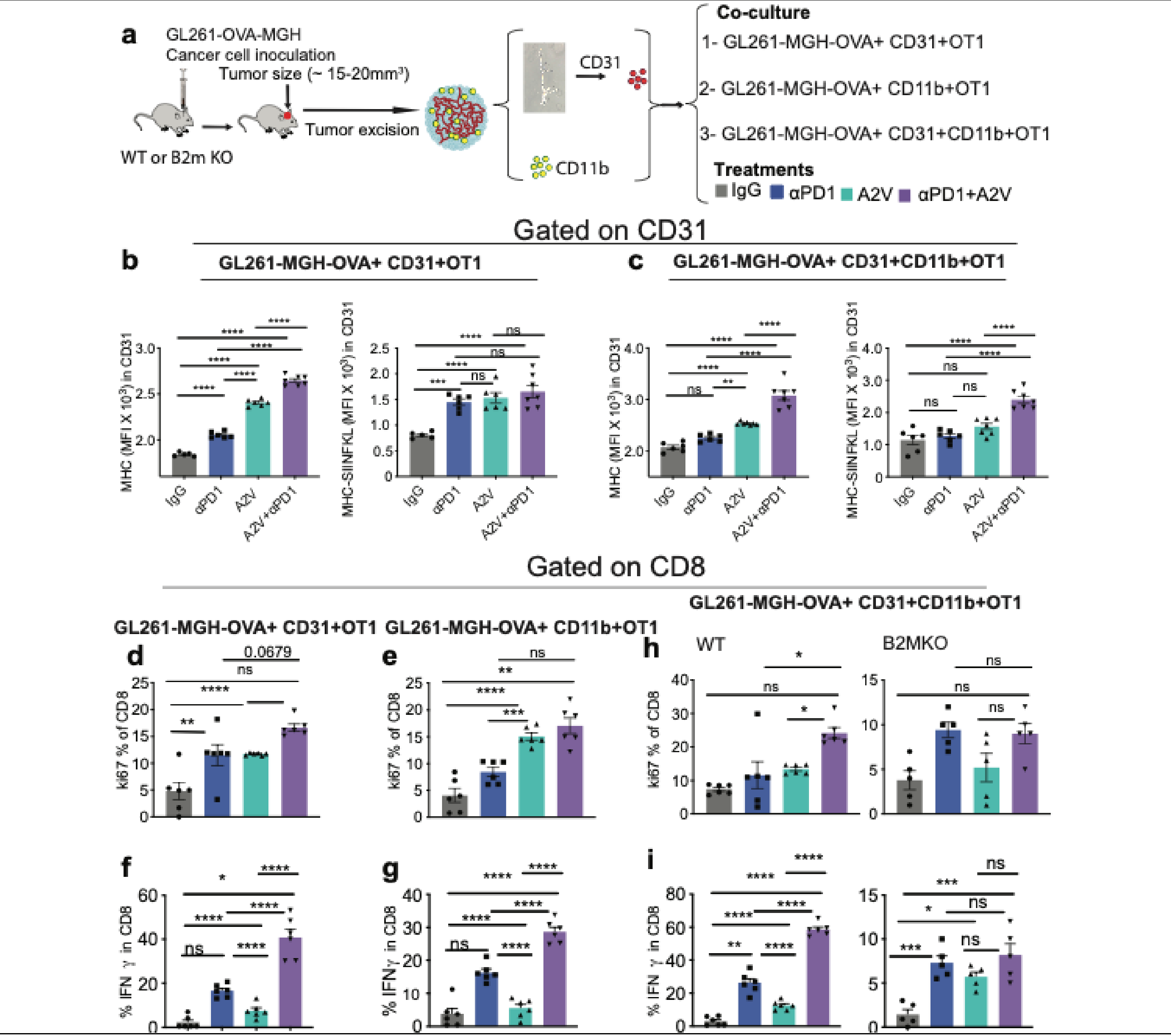


Supplemental Figure 8. ECs (non-professional antigen presenting cells) and classical antigen presenting cells in GBM are reprogrammed to present antigen to CD8 T cells after αPD1+A2V therapy. **a**) Schematic representation of *in vitro* co-culture assay. GL261-SIINFEKL-MGH cells are orthotopically implanted in WT or *B2m* KO mice. Tumors are excised when they reach a size of 15-20 mm$^3$. To enrich for CD11b cells, tumors are dissociated mechanically or with accutase ® into single cell suspension. Using Stem cell ® positive selection kit with a slight protocol modification (two round selection), CD11b cells are enriched at 95%. To enrich EC, whole blood vessels are harvested using ficoll gradient separation and then dissociated into single cells using 5 min accutase ® exposure at room temperature. WT CD8 T cells and OTI cells were enriched from the spleen using Stem cell technologies negative selection kit to a purity of 94-98%. Cells were co-cultured and treated as described in panel **a**. **b-c**) MFI (Mean Fluorescence Intensity) of MHC and MHC-SIINFEKL in (**b**) co-culture of cancer cells (GL261-SIINFEKL-MGH) with ECs (CD31) and antigen specific CD8 T cells (OTI) and (**c**) co culture of cancer cells (GL261-SIINFEKL-MGH) with ECs (CD31), classical antigen presenting cells (CD11b), and antigen specific CD8 T cells (OTI). **d-e**) Frequency of IFNγ+ cells in OT-I cells in the in (**d**) co-culture of cancer cells (GL261-SIINFEKL-MGH) with ECs (CD31) and antigen specific CD8 T cells (OTI) and (**e**) co-culture of cancer cells (GL261-SIINFEKL-MGH) with ECs (CD31), classical antigen presenting cells (CD11b), and antigen specific CD8 T cells (OTI). **f-i**) Comparison of the frequency of CD8 T cell effector function (IFNγ) in the co-culture of cancer cells (GL261-SIINFEKL-MGH) with ECs (CD31), classical antigen presenting cells (CD11b) harvested from WT versus *B2m*-KO mice.

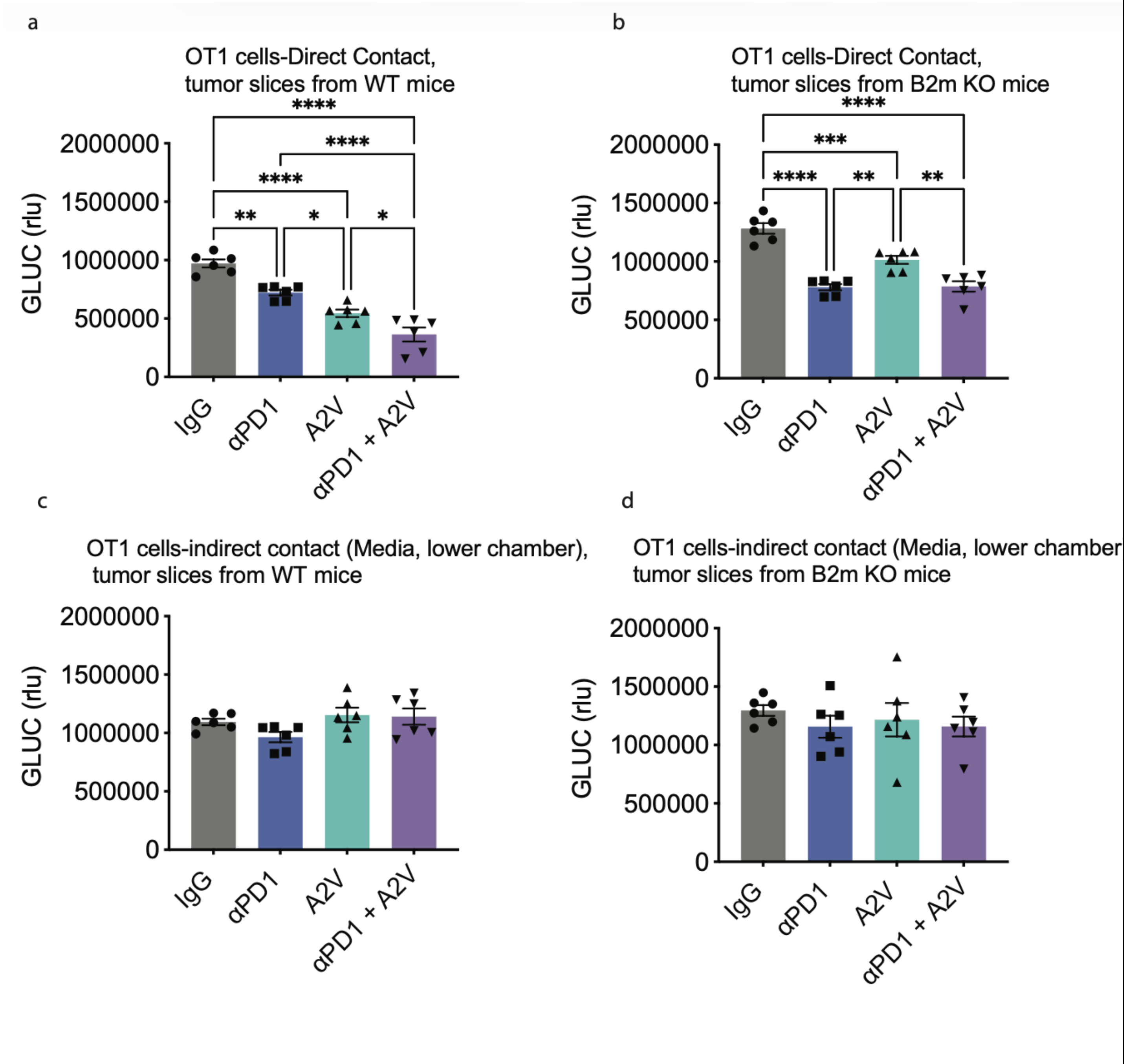

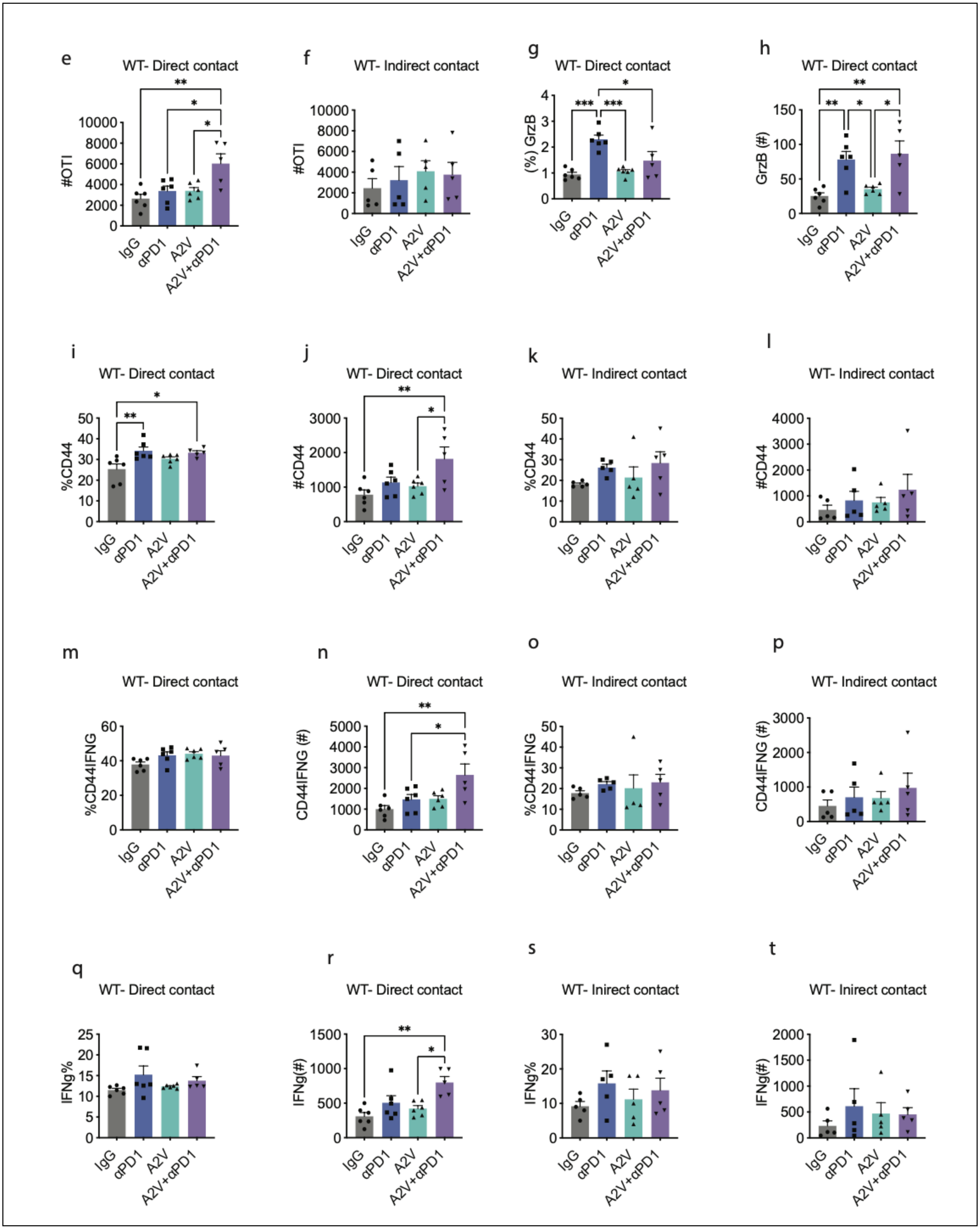
e
WT- Direct contact
#OTI
10000
8000
6000
4000
2000
0
**
*
*
IgG
αPD1
A2V
A2V+αPD1
f
WT- Indirect contact
#OTI
10000
8000
6000
4000
2000
0
IgG
αPD1
A2V
A2V+αPD1
g
WT- Direct contact
(%) GrzB
4
3
2
1
0
***
***
*
IgG
αPD1
A2V
A2V+αPD1
h
WT- Direct contact
GrzB (#)
150
100
50
0
**
**
*
*
IgG
αPD1
A2V
A2V+αPD1
i
WT- Direct contact
%CD44
50
40
30
20
10
0
*
**
IgG
αPD1
A2V
A2V+αPD1
j
WT- Direct contact
#CD44
3000
2000
1000
0
**
*
IgG
αPD1
A2V
A2V+αPD1
k
WT- Indirect contact
%CD44
50
40
30
20
10
0
IgG
αPD1
A2V
A2V+αPD1
l
WT- Indirect contact
#CD44
4000
3000
2000
1000
0
IgG
αPD1
A2V
A2V+αPD1
m
WT- Direct contact
%CD44IFNG
60
40
20
0
IgG
αPD1
A2V
A2V+αPD1
n
WT- Direct contact
CD44IFNG (#)
5000
4000
3000
2000
1000
0
**
*
IgG
αPD1
A2V
A2V+αPD1
o
WT- Indirect contact
%CD44IFNG
50
40
30
20
10
0
IgG
αPD1
A2V
A2V+αPD1
p
WT- Indirect contact
CD44IFNG (#)
3000
2000
1000
0
IgG
αPD1
A2V
A2V+αPD1
q
WT- Direct contact
IFNg%
25
20
15
10
5
0
IgG
αPD1
A2V
A2V+αPD1
r
WT- Direct contact
IFNg(#)
1500
1000
500
0
**
*
IgG
αPD1
A2V
A2V+αPD1
s
WT- Inirect contact
IFNg%
30
20
10
0
IgG
αPD1
A2V
A2V+αPD1
t
WT- Inirect contact
IFNg(#)
2000
1500
1000
500
0
IgG
αPD1
A2V
A2V+αPD1

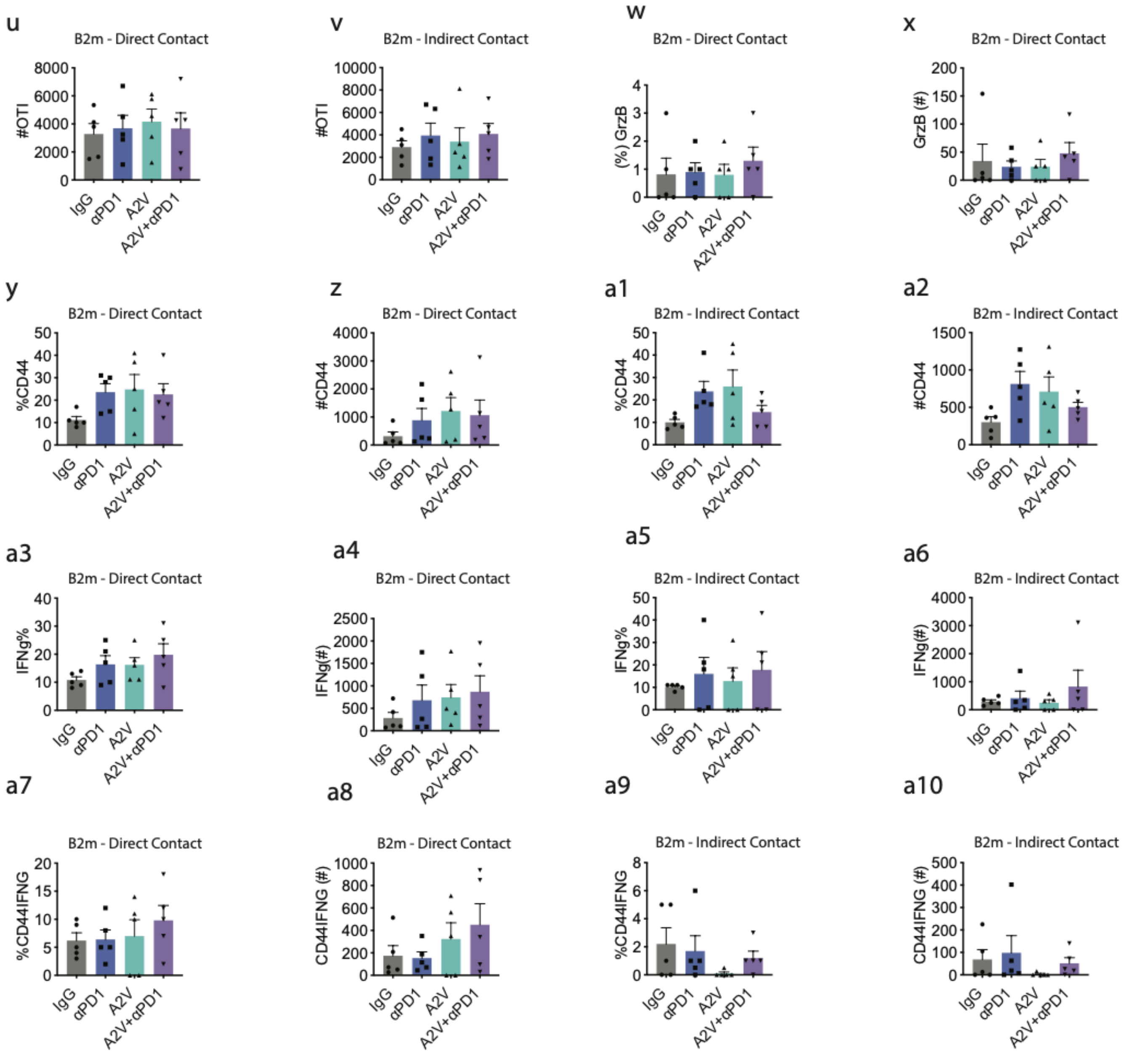


Supplemental Figure 9. Transwell cultures were set up such that brain tumor slices (grown in WT or *B2m* KO hosts) were cultured on the top well with OT-1 added either to the top (direct contact) or bottom (indirect contract) wells. Tumor volume measured by Gluc when OT-1 T cells are in direct contact with wildtype tumor slices grown in (a) WT or (b) *B2m* deficient mice. Tumor volume measured by Gluc when OT-1 T cells are indirectly cultured with tumor slices from WT hosts (c) or *B2m* deficient hosts (d). OT-1 T cell phenotype from cells in direct or indirect contact with tumor slice cultures grown in WT or *B2m* KO hosts (e-a10).

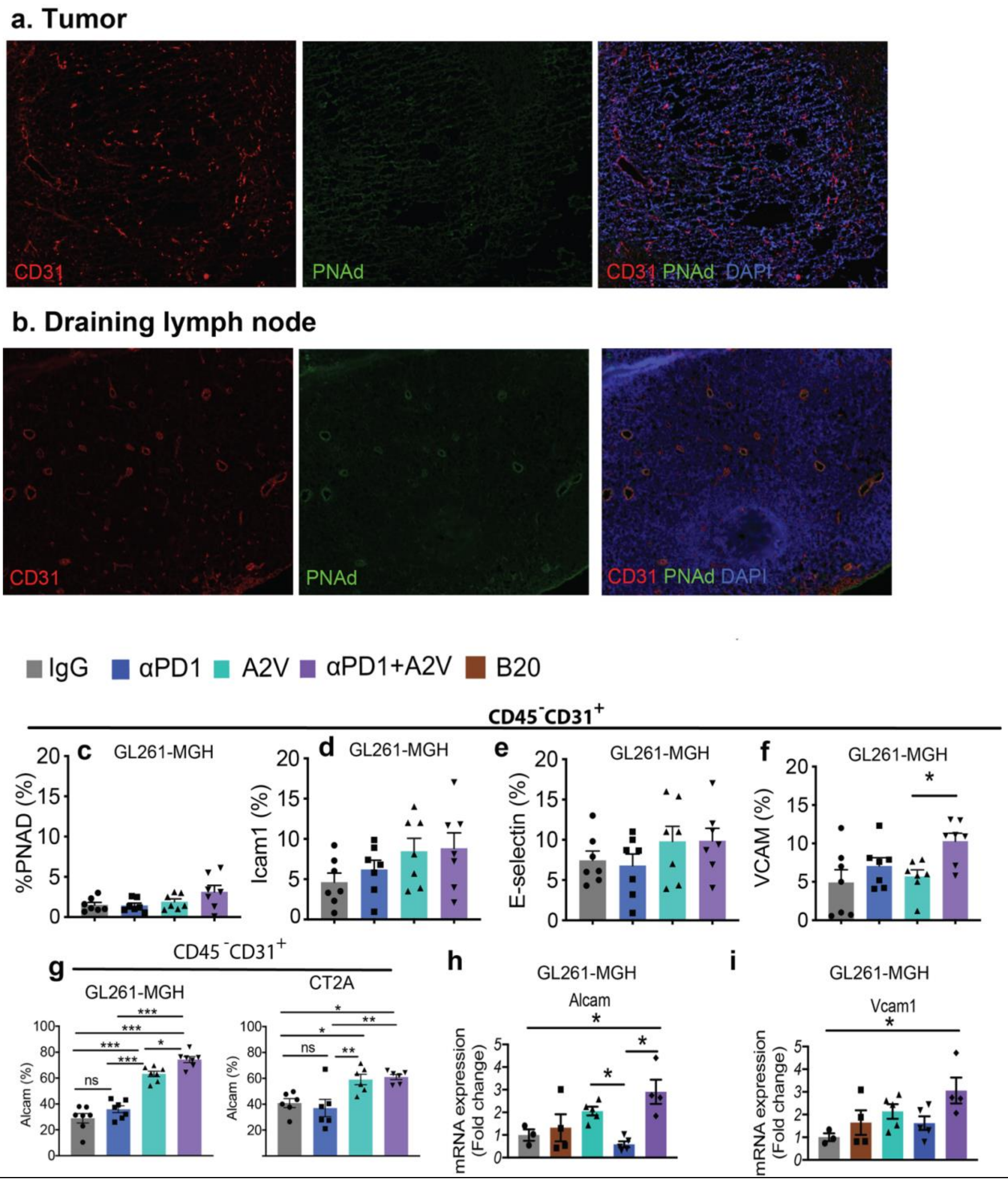


Supplemental Figure 10. Expression of PNAd in draining lymph node (dLN) versus GBMs. a) GBM and b) draining lymph node, fixed-frozen slides were co-stained for CD31 (red) and PNAd (green). **c-i**) Measured level of protein expression (**c**) PNAD $p$=0.0804, (**d**) ICAM $p$=0.1822, (**e**) E-selectin $p$=0.3689, (**f**) VCAM1 $p$=0.0217, and (**g**) ALCAM $p$=0.0011 using flow cytometry after IgG (n=7), αPD1 (n=7), A2V (n=7), αPD1+A2V (n=7). **h-i**) mRNA levels of *Alcam* (**h**) and *Vcam1* (**i**) in endothelial cells using QPCR, after IgG (n=3), αPD1 (n=5), B20 (n=4), A2V (n=5), αPD1+A2V (n=4), P<0.001. Data presented are mean ± SEM. Multiple comparisons using the Tukey adjustment. For between-group analysis post-Tukey, Statistical significance is shown as *$p$ < 0.05, **$p$ < 0.01, ***$p$ < 0.001, ****$p$ < 0.0001.

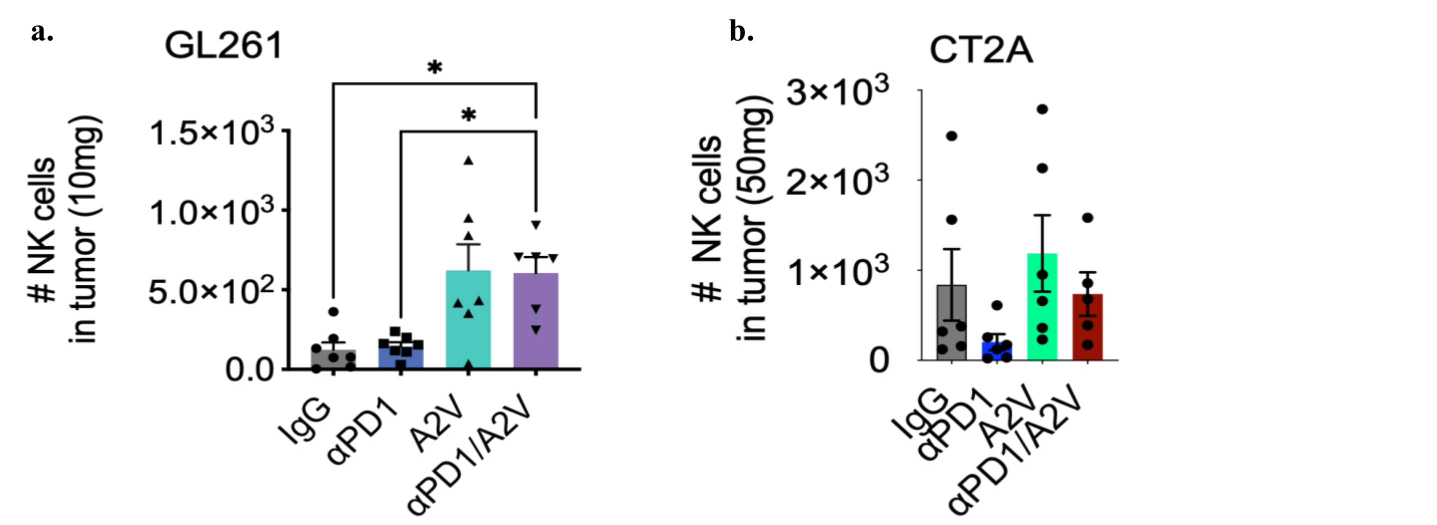


Supplemental Figure 11. Flow cytometric analysis of NK cells in GL261 (a) or CT2A (b) tumors following IgG, aPD1, A2V, or combined aPD1/A2V treatment.

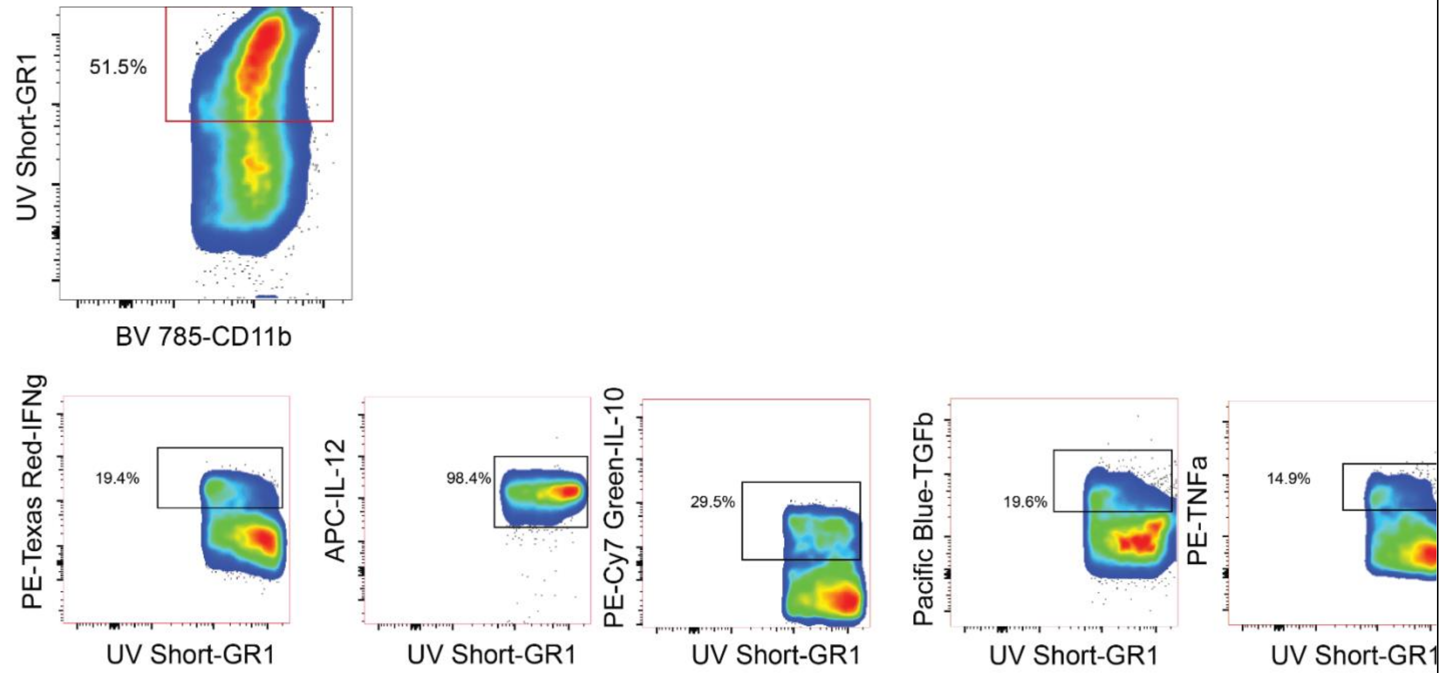


Supplemental Figure 12. Gating strategy for CD11b cells and classical MDSC, marked by GR1. GR1+ cells produce a range of inflammatory (IFNγ, IL12, and GR1) and anti-inflammatory cytokines (IL-10 and TGFβ) in GBM.

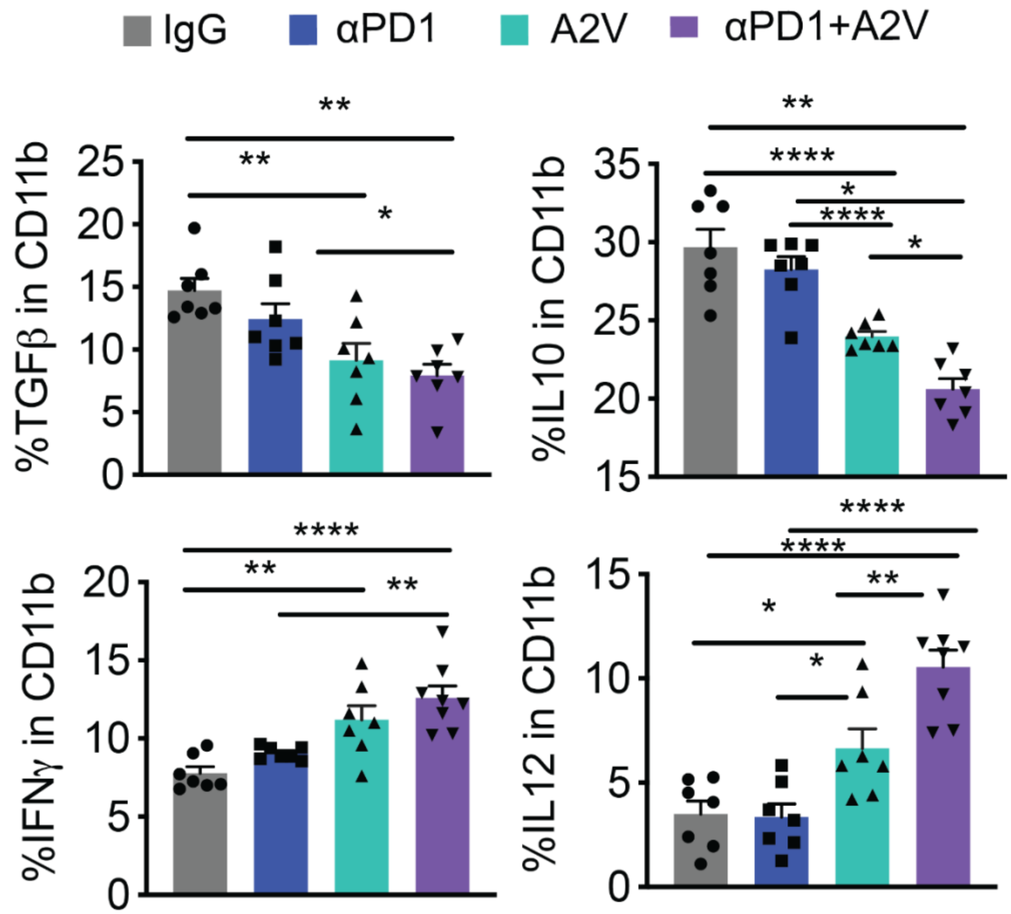


Supplemental Figure 13. GBM bearing GL261-MGH mice treated with αPD1+A2V have reduced production of anti-inflammatory cytokine (TGFβ [IgG (n=7), αPD1(n=7), A2V (n=7), αPD1+A2V (n=7) $p = 0.001$] (left) and IL10 [IgG (n=7), αPD1(n=7), A2V (n=7), αPD1+A2V (n=7) $p < 0.0001$] (right), first row) and enhanced production of inflammatory cytokine (IFNγ [IgG (n=7), αPD1(n=7), A2V (n=7), αPD1+A2V (n=7) $p < 0.0001$] (left), and IL12 [IgG (n=7), αPD1(n=7), A2V (n=7), αPD1+A2V (n=7) $p < 0.0001$] (right), second row) compared by ordinary one-way ANOVA test and corrected for multiple comparisons using the Tukey adjustment. For between group analysis post-Tukey, Statistical significance is shown as *$p < 0.05$, **$p < 0.01$, ***$p < 0.001$, ****$p < 0.0001$.

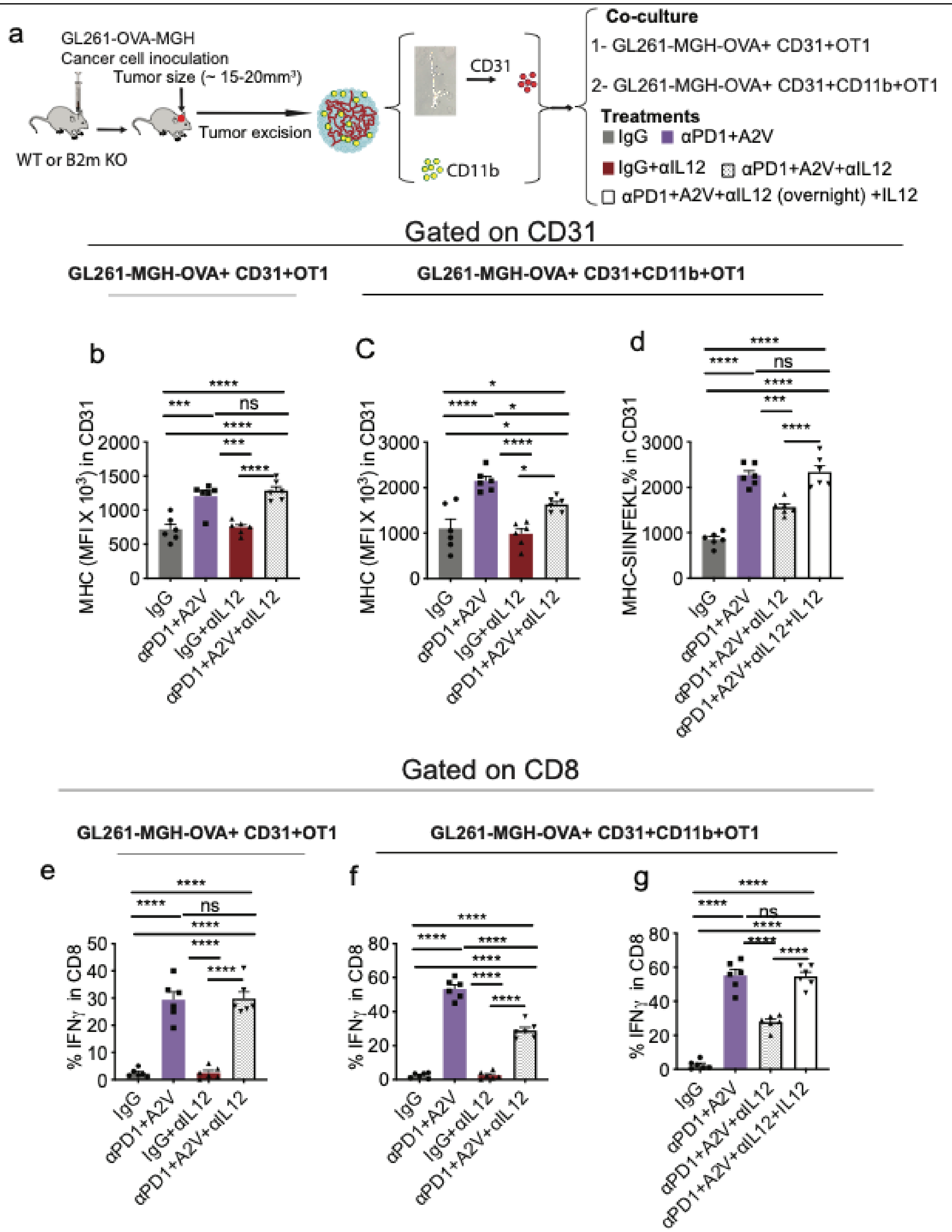


Supplemental Figure 14. IL-12 production by CD11b cells after αPD1+A2V therapy contributes to CTL's IFNγ production to restore antitumor immunity. **a**) Schematic representation of *in vitro* co-culture assay. GL261-SIINFEKL-MGH cells were orthotopically implanted in mice. Tumors were excised and processed as and co-cultures were treated as shown. **b-c**) Mean Fluorescence Intensity (MFI)of MHCI in ECs: **b**) co-culture of cancer cells (GL261-SIINFEKL-MGH) with ECs (CD31) and antigen specific CD8 T cells (OTI) and (**c**) MFI of MHCI and (**d**) MHCI-SIINFEKL co-culture of cancer cells (GL261-SIINFEKL-MGH) with ECs (CD31), classical antigen presenting cells (CD11b), and antigen specific CD8 T cells (OTI).

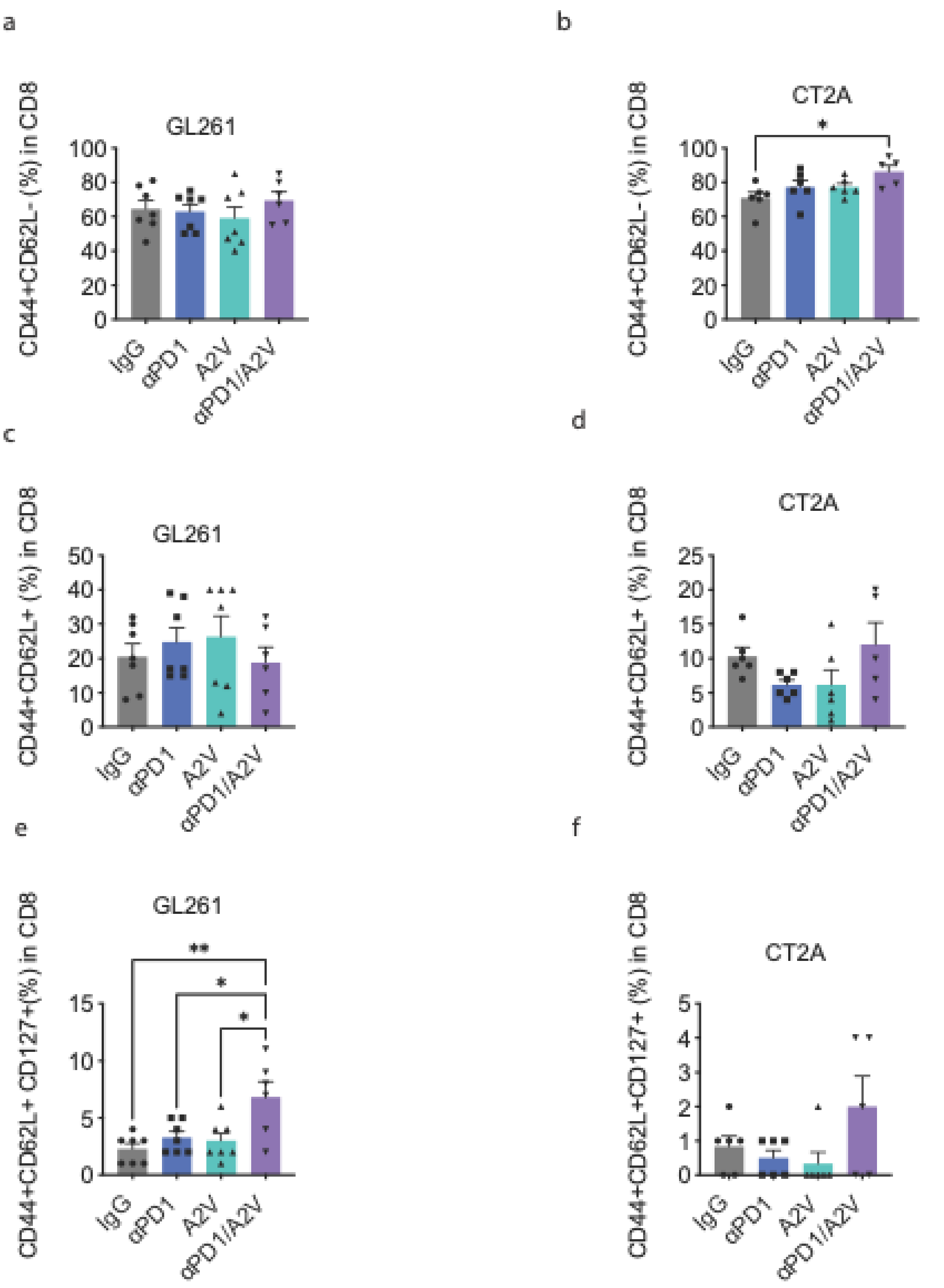


Supplemental Figure 15. Memory Markers on T cells from GL261 or CT2A following IgG, aPD1, A2V, or combination treatment. (a) Expression of CD44+ CD62L- CD8 T cells in GL261 or (b) CT2a. (c) Expression of CD44+ CD62L+ CD8 T cells in GL261 or (d) CT2a. (e) Expression of CD44+ CD62L+ CD127+ CD8 T cells in GL261 or (f) CT2a.
